# Investigating the Functional Consequence of White Matter Damage: An Automatic Pipeline to Create Longitudinal Disconnection Tractograms

**DOI:** 10.1101/140137

**Authors:** Kesshi Jordan, Anisha Keshavan, Eduardo Caverzasi, Joseph Osorio, Nico Papinutto, Bagrat Amirbekian, Mitchel S. Berger, Roland G. Henry

## Abstract

Neurosurgical resection is one of the few opportunities researchers have to image the human brain both prior to and following focal damage. One of the challenges associated with studying brains undergoing surgical resection is that they often do not fit the brain templates most image-processing methodologies are based on, so manual intervention is required to reconcile the pathology and the most extreme cases must be excluded. Manual intervention requires significant time investment and introduces reproducibility concerns. We propose an automatic longitudinal pipeline based on High Angular Resolution Diffusion Imaging acquisitions to facilitate a Pathway Lesion Symptom Mapping analysis relating focal white matter injury to functional deficits. This two-part approach includes (i) automatic segmentation of focal white matter injury from anisotropic power differences, and (ii) modeling disconnection using tractography on the single-subject level, which specifically identifies the disconnections associated with focal white matter damage. The advantages of this approach stem from (1) objective and automatic lesion segmentation and tractogram generation, (2) objective and precise segmentation of affected tissue likely to be associated with damage to long-range white matter pathways (defined by anisotropic power), (3) good performance even in the cases of anatomical distortions by use of nonlinear tensor-based registration in the patient space, which aligns images using white matter contrast. Mapping a system as variable and complex as the human brain requires sample sizes much larger than the current technology can support. This pipeline can be used to execute large-scale, sufficiently powered analyses by meeting the need for an automatic approach to objectively quantify white matter disconnection.

Abbreviations

DTI: Diffusion Tensor Imaging
IOS: Intra-Operative Stimulation
VLSM: Voxel-Based Lesion-Symptom Mapping
MD: mean diffusivity
FA: fractional anisotropy
B0: minimally diffusion-weighted image
AP: anisotropic power
ASAP: automatic segmentation of anisotropic power changes
HARDI: High Angular Resolution Diffusion Imaging
MRI: Magnetic Resonance Imaging
FSL: FMRIB Software Library
Dipy: Diffusion Imaging in Python
APM: Anisotropic Power Map was calculated
DTI-TK: Diffusion Tensor Imaging ToolKit
TFCE: Threshold-Free-Cluster-Enhancement
ROI: Region of Interest
CCI: Cluster Confidence Index
AF: arcuate Fascicle
SLF II and SLF III: components 2 and 3 of the SLF
SLF-tp: temporo-parietal component of the SLF
IFOF: inferior fronto-occipital Fascicle
UF: uncinate Fascicle
ILF: inferior longitudinal Fascicle
Md-LF: middle longitudinal Fascicle
CST: corticospinal tract
OR: optic radiation
QC: quality-control

**Funding:** This work was supported by the National Institutes of Health [5R01NS066654-05]; KJ was supported by the Department of Defense (DoD) [National Defense Science & Engineering Graduate Fellowship (NDSEG) Program].

## 1 Introduction

### 1.1 Background

The relationship between injury to the brain and behavioral changes has been a keystone of human neuroscience research for well over a century (Berti, Garbarini, and Neppi-Modona 2015; Geschwind 1965; Bates et al. 2003). Studying the spatial location of white matter injury, however, is often insufficient to predict functional outcome because disconnection of long myelinated pathways can have widespread effects on systems employing the disrupted circuit, manifesting in apparently inconsistent deficits (Geschwind 1965; Geschwind 1965). Diffusion Tensor Imaging (DTI) (Basser, Mattiello, and LeBihan 1994; Le Bihan et al. 2001; Pierpaoli et al. 1996; Pajevic and Pierpaoli 1999) and the more complex models that followed (Tuch 2004; Tournier et al. 2004; Jensen et al. 2005; Assaf and Basser 2005; Jeurissen et al. 2014; Tuch et al. 2002; Hess et al. 2006) have enabled researchers to investigate disconnection syndromes with improved specificity to white matter structure in a variety of clinical applications (Kumar et al. 2016; Kumar et al. 2016; Yogarajah et al. 2009; Chen et al. 2009; Glenn et al. 2003; Mandelli et al. 2014; Jang 2013) using fiber tracking, also called tractography, to infer connectivity (Mori and van Zijl 2002). Tractography has been used extensively to study the clinical consequence of white matter disconnection by modeling the white matter connectivity on a single-subject level and associating damage to white matter structures with functional deficits (Caverzasi et al. 2016; Duffau 2008; Kim and Jang 2012), but there is a need for an objective longitudinal approach to studying focal white matter damage and modeling the downstream effects caused by the disconnection to guide the clinical translation of these methods currently underway.

### 1.2 Clinical Translations

Elucidating the functional impact of new focal or progressive injury to the brain’s intricate communication network is relevant to a wide range of clinical disorders. This relationship has typically been investigated by either registering an estimate of the injurious region of interest to a template of putative fiber bundles (volume-based method), or by modeling those fiber bundles in each patient using tractography (connectivity-based method). Tractography reconstruction of white matter structures for pre-surgical planning and guidance of intra-operative cortical and subcortical stimulation mapping has emerged as a major clinical translation of diffusion MRI applications. Tractography methods have shown great potential to improve clinical outcome in neurosurgery when used in tandem with Intra-Operative Stimulation (IOS) (Wu et al. 2007; Leclercq et al. 2010; Henry et al. 2004; Lehéricy et al. 2007; Lehéricy et al. 2014; Berman et al. 2007; Bucci et al. 2013; Mandelli et al. 2014; Caverzasi et al. 2016; Berman 2013). However, tractography is an inexact science that is highly dependent on implementation choices (Chamberland et al. 2014; Fillard et al. 2011; Crettenand et al. 2006; Neher et al. 2015) and often a human operator, exacerbating well-documented reproducibility concerns (Wakana et al. 2007; Berman 2013) so translation to the clinic requires extensive validation (Kinoshita et al. 2005; Duffau 2014). The added complexity of pathology introduces additional errors, which can vary with tractography algorithm choices and underlying anatomy (Golby et al. 2011; Chamberland et al. 2014). Establishing robust relationships between deficits and damage patterns using reproducible, operator-independent methods is essential to being able to guide neurosurgical intervention at the single-subject level.

### 1.3 Volume- and Connectivity-Based Approaches to Lesion Studies

There are several ways to evaluate the impact of *focal white matter damage* on fiber pathways that reflect different levels of efficiency and accuracy. The challenges are (i) segmentation of the focal injury and (ii) association of this focal injury with fiber pathways. The simplest approach is to localize damage on an anatomical image and infer what structures were likely to have been damaged either using a white matter atlas (Ius et al. 2011), healthy controls (Meyer et al. 2016), or judgment by an expert (ex: neuroradiologist). A widely used example of this approach is called Voxel-Based Lesion-Symptom Mapping (VLSM), which has been used to great effect in leveraging modern imaging to conduct these lesion studies (Bates et al. 2003; Kinoshita et al. 2016; Campana, Caltagirone, and Marangolo 2015; Almairac et al. 2014).

While volume-based methods can be an efficient and straightforward approach using conventional imaging, the brains subjected to pathological or neurosurgical damage often will present unique distortions resulting in misregistration to the template. Current methods used to address some of these concerns (Brett, Leff, and Ashburner 2000; Ashburner and Friston 2005; Andersen, Rapcsak, and Beeson 2010) enable registration of brains with distorted anatomy, but are still prohibitively labor-intensive for large studies and introduce reproducibility concerns inherent to manual tracing (Fiez, Damasio, and Grabowski 2000). Alternative methods are being developed to address these challenges (Werner et al. 2016), but many of these approaches are designed for stroke applications and may not allow wider applicability for cases with increased heterogeneity and complexity, like brain tumors (Gordillo, Montseny, and Sobrevilla 2013; Menze et al. 2015; Sanjuán et al. 2013).

A second approach to evaluate focal white matter damage is to reconstruct fascicles using tractography and inferring damage to the structure using properties of the segmented volume (e.g. stroke), or apparent displacement/destruction of the reconstructed model (e.g. tumor). In the neurosurgical example, these studies can be conducted longitudinally because the state of the brain’s white matter prior to the injury can be acquired. One way to execute this type of study is to model the fascicle pre-operatively and infer damage using a segmentation of the post-operative resection cavity. An alternative approach is to reconstruct the fascicle model using tractography at two time points (pre- and post-surgery) and judge how the integrity of the fascicle changed based on the track model (Caverzasi et al. 2016). These approaches have enabled researchers to make great strides in understanding the relationship between focal damage and functional deficits, but they are limited by their dependence on high-fidelity registration and/or subjective intervention in the cases of complex pathology, like tumors.

### 1.4 Technical Challenges: Tumor Resection Studies

Executing group imaging studies in the field of brain tumor surgery is particularly difficult (Brennan and Holodny 2016). The heterogeneity of brain tumor pathology, often presenting with solid, infiltrative components and perilesional vasogenic edema, can change the diffusion properties in tissues-of-interest. Moreover, tumor growth can displace or disrupt pathways (Field et al. 2004). Tissue can shift considerably during surgery (Nimsky et al. 2005), so spatial relationships between pre-, intra-, and post-operative imaging may not be maintained. Imaging post-operatively to evaluate the effect of surgical intervention has unique challenges, as well. The presence of intracranial blood products, air, and metallic implants used on the surface of the skull in craniotomies can cause imaging artifacts in some modalities. Taken together, these complications cause many of the automatic computational tools used to extract salient features from large neuroimaging datasets to fail, requiring significant human time investment and/or exclusion of cases. As a result, the typical association study between tractography models and deficits is underpowered and the results are often operator-dependent. There is a need for an automatic method to objectively quantify focal white matter injury.

### 1.5 Proposed Method: Automatic Pipeline to Create Disconnection Tractograms

In this manuscript, we present an automatic pipeline for comparing diffusion MRI volumes and identifying spatially coherent focal white matter disconnection. This pipeline was developed based on a tumor cohort undergoing surgical resection, but the method could be extended to study longitudinal focal white matter changes in many other contexts.

The proposed method (ASAP-tractograms) has two essential components to address the challenges outlined, above. The first component is an automatic segmentation of anisotropic power (AP) changes (ASAP), which identifies tissue likely to be associated with damage to long-range pathways based on diffusion properties. This approach bypasses the challenges inherent to automatic segmentation and classification of various lesion components to solely focus on focal white matter damage. The second component involves modeling disconnection using tractography on the single-subject level with High Angular Resolution Diffusion Imaging (HARDI)-based tractograms. This approach enables a comparison between subjects that is based on individual characterization of damage to long-range pathways, so comparisons between patients are made based directly on white matter structures. The advantages of this approach stem from (1) objective and automatic lesion segmentation and tractogram generation, (2) objective and precise segmentation of affected tissue likely to be associated with damage to long-range white matter pathways (defined by AP), (3) good performance even in the cases of anatomical distortions by use of diffusion MRI based registration in the patient space, which aligns images using white matter contrast. To evaluate this method against an established approach, we re-analyze a previously published study (Caverzasi et al. 2016) relating white matter damage to functional deficits in a neurosurgical cohort and compare the ASAP disconnection tractograms to the subjective damage ratings of each fascicle established by a consensus of two trained neuro-radiologists.

## 2 Material and Methods

### 2.1 Subjects

The ASAP-Tractogram pipeline was evaluated on pre and post-surgical resection MRI data from thirty-five subjects with glioma brain tumors. Research was performed in compliance with the Code of Ethics of the World Medical Association (Declaration of Helsinki) and the standards established by our institution. The Committee on Human Research at the University of California, San Francisco, approved the study protocol. Written informed consent was obtained from all study participants. Each subject underwent a standard pre-surgical Magnetic Resonance Imaging (MRI) scanning protocol including a High Angular Resolution Diffusion Imaging sequence. This cohort is the same used in a previous publication; details can be found in (Caverzasi et al. 2016). Of the 35 patients in this cohort, 6 were excluded for acquisition-related reasons (3 scan-rescan dates greater than a week apart, 2 sequence mismatches, 1 had a major artifact obscuring a large portion of the frontal lobe), leaving 29 patients to be analyzed.

### 2.2 Diffusion Preprocessing

Diffusion preprocessing was applied to both pre- and post-surgery HARDI datasets in parallel, as shown in Supplementary Figure 1. The datasets were first corrected for motion and eddy current distortion using the FMRIB Software Library (FSL) (Jenkinson et al. 2012) and the gradient table rotated, accordingly (Leemans and Jones 2009). A tensor model was fit to the corrected HARDI data using the open-source package Diffusion Imaging in Python (Dipy) (Garyfallidis et al. 2014) and the resulting parameters used to calculate tensor metrics. An Anisotropic Power Map was calculated (APM) (Flavio Dell’Acqua and Simmons 2014) using the Q-ball model (Descoteaux et al. 2007) implemented in Dipy, which should represent white matter better than Fractional Anisotropy, especially in crossing regions (Flavio Dell’Acqua and Simmons 2014). The brain mask calculated from the B0 image (Smite 2002) was applied to all images.

### 2.3 Tensor-Based Longitudinal Alignment

Rigid and non-linear tensor registration was performed to align white matter structures at the two time points using the Diffusion Tensor Imaging ToolKit (DTI-TK) (Zhang et al. 2006). These images were subtracted pre-surgery minus post-surgery to produce peri-surgical difference maps, shown in Supplementary Figure 2. To segment the resection cavity, the flowchart shown in Figure 1 was applied. The registered pre- and post-surgical Anisotropic Power Maps were subtracted and spatially clustered using FSL’s Threshold-Free-Cluster-Enhancement (TFCE) (Smith and Nichols 2009; Woolrich et al. 2009; Smith et al. 2004; Jenkinson et al. 2012) to boost the signal of spatially coherent decrease in AP. This made it possible to isolate the ASAP ROI using simple thresholds. These TFCE maps were thresholded at the 99th percentile, which was sufficient to isolate the AP change associated with the resection in most patients as a region of interest (ROI), but additional noise from tissue shifting necessitated the additional requirements of a hard threshold TFCE-boosted intensity >500 and a cluster size of >5. This results of 29 patients were quality-controlled using the Mindcontrol platform (Keshavan et al. 2017) for accurate representation of post-surgical cavity.

**Figure 1:**
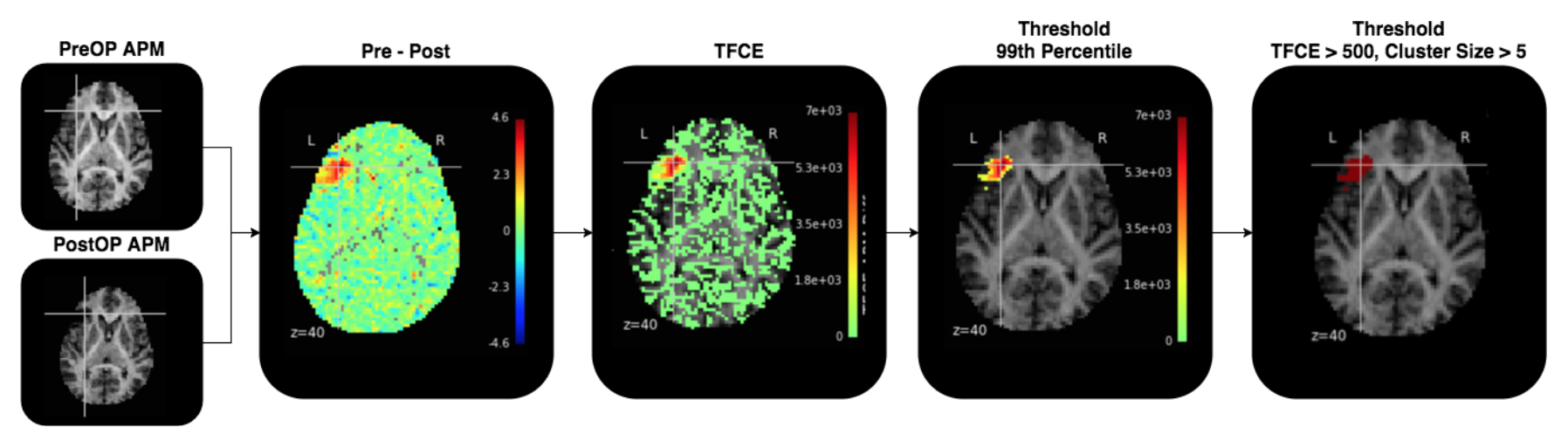
The pre- and the post-surgical Anisotropic Power Maps are subtracted to produce the Pre - Post image. This image is spatially clustered using FSL’s threshold-free cluster enhancement to boost the value of spatially coherent decreases in AP, resulting in the TFCE image. Several thresholds were applied (99^th^ percentile, TFCE-boosted intensity >500, cluster size >5) to isolate the ASAP ROI.

### 2.4 Tractography

The whole-brain streamline dataset was targeted using the ASAP ROI. The resulting distribution of streamlines represents likely pathways of major underlying white matter structures at risk of being damaged by the surgical resection. The pathways with higher confidence have many streamlines following roughly the same trajectory. This confidence can be estimated using the Cluster Confidence Index (CCI) (Jordan et al. Accepted). To eliminate low-confidence pathways that make results noisy and difficult-to-interpret, any streamline shorter than 40mm or with CCI <1 (calculated using default parameters: k=1, theta=5, subsamp=8) was excluded from figures and analysis (Supplementary Figure 3).

Any tractography method can be applied to produce a prediction of the white matter structures disrupted by the focal injury. For the demonstration detailed by this manuscript, the pre-surgical whole brain white matter was seeded (AP >4) and tracked using residual bootstrap probabilistic q-ball tractography (Berman et al. 2008) with the parameters described in (Caverzasi et al. 2016) (Tristán-Vega, Westin, and Aja-Fernández 2009; Tristán-Vega and Aja-Fernández 2010; Tristán-Vega and Westin 2011; Bucci et al. 2013).

### 2.5 Software Implementation

This pipeline was built and run using the Nipype software package (Gorgolewski et al. 2011). The code is open-source and can be downloaded from GitHub (https://github.com/kesshijordan).

### 2.6 Comparison to Manual Analysis

The results of this pipeline were compared to those generated by (Caverzasi et al. 2016). In (Caverzasi et al. 2016), the following white matter bundles were segmented independently by two neuroradiologists: arcuate Fascicle (AF), components 2 and 3 of the SLF (SLF II and SLF III), temporo-parietal component of the SLF (SLF-tp), inferior fronto-occipital Fascicle (IFOF), uncinate Fascicle (UF), inferior longitudinal Fascicle (ILF), middle longitudinal Fascicle (Md-LF), according to methods outlined in their cited paper (Caverzasi et al. 2016; Catani, Jones, and ffytche 2004; Caverzasi et al. 2014; Makris et al. 2012), along with the corticospinal tract (CST) and optic radiation (OR) (Bucci et al. 2013) (Hofer 2010). The same two neuroradiologists independently rated the degree to which each of these fascicles had been disrupted both pre- and post-surgery as unchanged (0), displaced but otherwise normal appearing (1), partially interrupted (2), or completely interrupted (3). These categories were reduced to affected (0 or 1) and not affected (2 or 3). A pre- to post-surgical change from 1 to 2 or from 2 to 3 should be reflected in the results of the difference pipeline because either of those transformations indicate removal of tissue associated with a disconnection.

## 3 Results

Disconnection tractograms are shown for the entire cohort that passed quality-control (QC) (26/29 subjects) in Figure 2.

**Figure 2:**
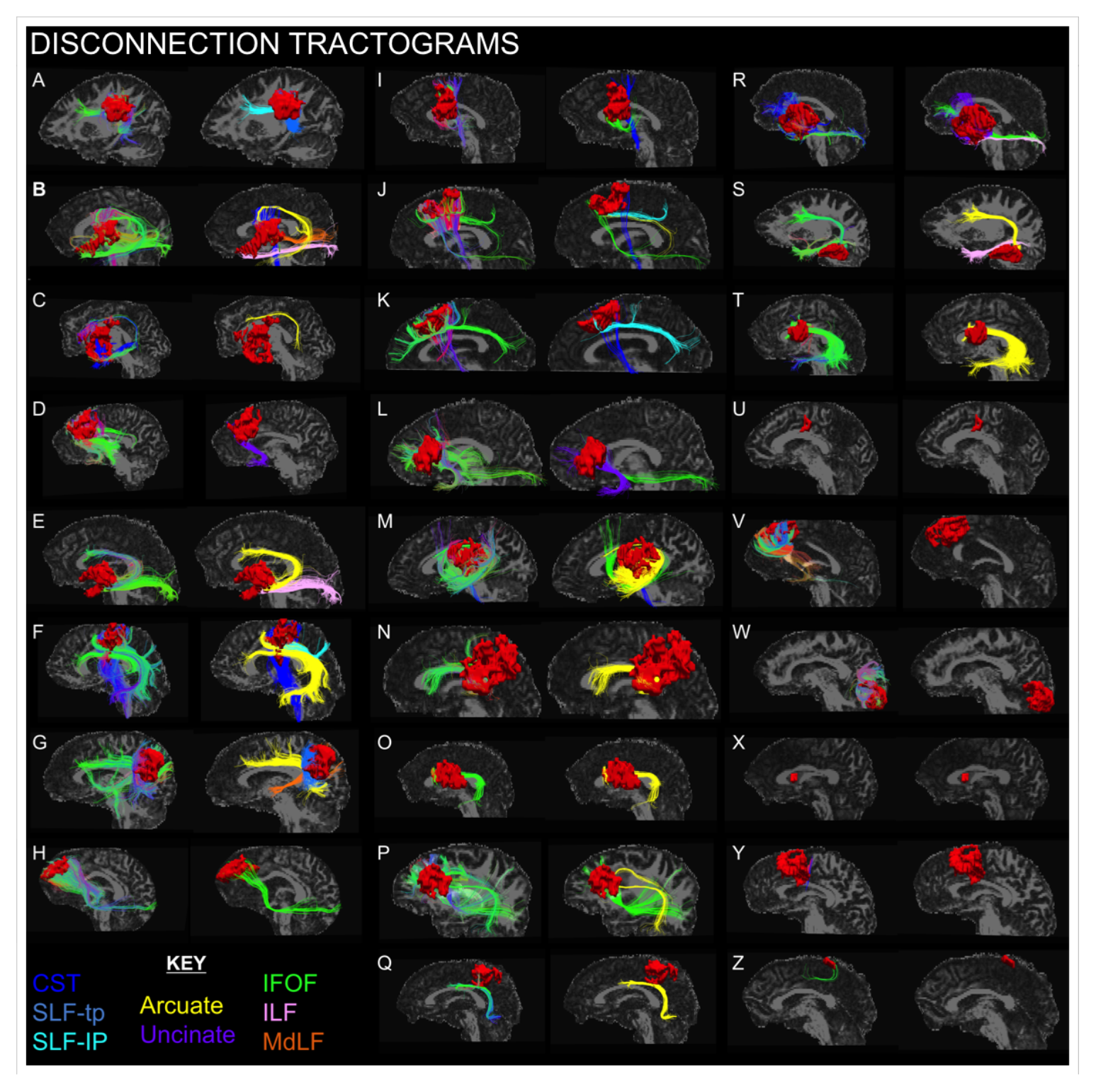
Shows the Disconnection Tractograms and identified tracts in this cohort that passed quality control. The left image shows the resection ROI (red) with the raw tractography output, colored by standard direction. The right image shows the tractography output segmented by a human operator to reflect fascicle membership. The left image for each patient shows the automatically-generated connectivity of the ASAP ROI (CCI >1), as described in the methods, colored by standard orientation. In the standard orientation color-scheme, each streamline is colored according to the dominant trajectory, as seen on the traditional FA color map (red=right/left; blue=superior/inferior; green=anterior/posterior). The right image shows any fascicles-of-interest colored according to the provided key [CST=cortico-spinal track (dark blue); SLF-tp=superior longitudinal fascicle temporal-parietal component (medium blue); SLF-IP=superior longitudinal fascicle infra-parietal component (light blue); Arcuate=arcuate fascicle (yellow); Uncinate=uncinate fascicle (purple); IFOF=inferior fronto-occipital fascicle (green); ILF=inferior longitudinal fascicle (pink); MdLF=Medial longitudinal fascicle (orange)]. Of the 29 patients that met inclusion criteria, 3 did not pass quality control: one patient had a bad registration between the pre- and post-surgical images and 2 had false-positive ASAP ROI’s present. In the 3 cases investigated for anomalous large regions of Anisotropic Power increase post-surgery, other imaging contrasts indicated postoperative blood products and/or pneumocephalus colocalizing with the region.

### 3.1 Evaluation of Performance: Connectivity of ASAP ROI

The streamline output of the difference pipeline clearly shows the white matter structures hypothesized to have been disrupted by the surgery (Figure 3). These sub-bundles are recognizable by shape and their pathway through the brain. For the purposes of demonstrating this using a two-dimensional figure, the streamline output for each case has been colored by a researcher with experience in fascicle modeling to highlight the bundles associated with motor, optic, or language that are routinely modeled for pre-surgical planning research at UCSF (Figure 2). These illustrative cases are compared to the subjective rating conducted by (Caverzasi et al. 2016). This study manually segmented fascicle models of bundles pre- and post-surgery, rating each subjectively on a scale from 0 to 3 (0=unaffected, 1=infiltrated/displaced, 2=partially destroyed, 3=completely destroyed). The difference pipeline shows disconnections in tractography models, so we would expect bundles output by the difference pipeline to reflect rating changes from 0/1→2 or 0/1/2→3.

**Figure 3:**
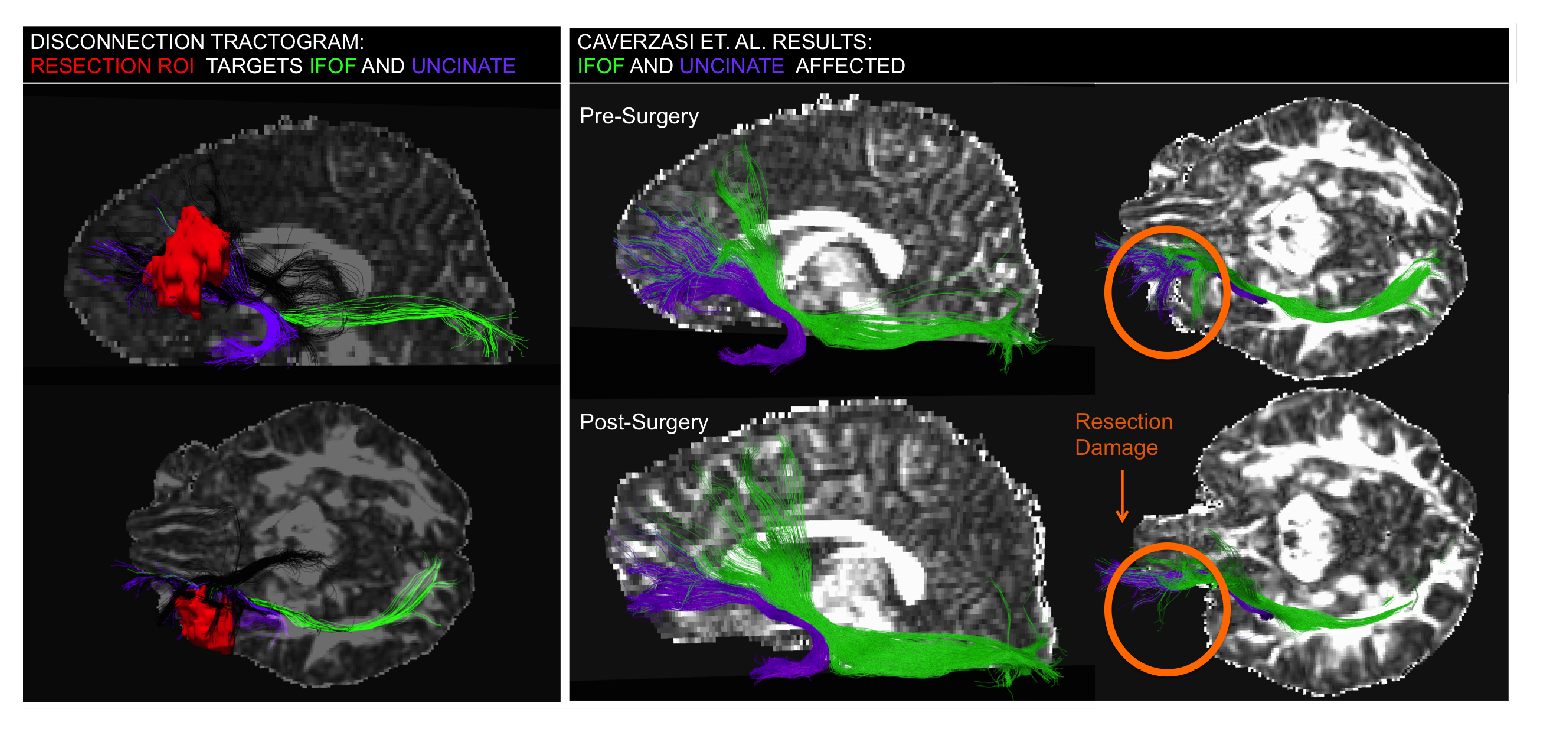
Comparing Difference Pipeline to Pre-/Post-Surgery Anatomically Constrained Fascicle Models LEFT: The Difference Pipeline Results suggest that the resection ROI (red) disconnected segments of the Inferior Fronto-Occipital (green) and the Uncinate (purple) Fasciculi. The RIGHT panel shows the manual/subjective comparison (Caverzasi 2016) study results pre (LEFT) and post (RIGHT) surgery in the transverse plane. The orange circle indicates the change pre- vs. post-surgery. The manual results indicated potential disruption of the SLFII, but the underlying white matter is intact so it is likely due to post-surgical tracking difficulty should not be included in the disconnection tractogram. Manual Results: IFOF (1→2), Uncinate (1→2), SLFII (2→3), SLFIII (2→2), Arcuate (1→1)

Figure 3 demonstrates the basic idea: the left panel shows the connectivity of the difference pipeline output ASAP ROI (colored red), as modeled by tractography. We can see a bundle that follows the path of the Uncinate Fascicle (purple) from the frontal lobe, hooking down and around into the anterior temporal lobe and another bundle that follows the path of the IFOF (green), from the frontal lobe through the external/extreme capsule and continuing posteriorly to terminate in the occipital lobe. The connectivity of the ASAP ROI suggests that portions of these two bundles were disconnected by the surgery. The right panel shows the pre- and post-surgery tractography models from which the damage rating was assigned. From these manual reconstructions, it appears that the lateral projections of both the UF and IFOF present in the pre-surgery model are missing from the post-surgery model. In this case, the subjective damage rating of the UF and IFOF increased from 1 (infiltrated/displaced) to 2 (partially destroyed), so the approaches reached the same conclusion with respect to these two structures. The subjective rating also scored the SLF II as increasing from a score of 2 to 3, but the resection cavity did not overlap with the missing streamlines from the SLF II (the underlying white matter appears intact), so this disagreement is not relevant.

### 3.2 Continuous Representation of White Matter Damage

One advantage of representing white matter damage continuously, as opposed to using a rating scale, is that some damage patterns cannot be described discretely. Figure 4 demonstrates this point. From the connectivity of the ASAP ROI (Figure 4 left panel), portions of the left Arcuate Fascicle (yellow) and left IFOF (green) can be segmented. This agrees with the manual reconstructions of the pre- and post-surgery Arcuate and IFOF (Figure 4 right panel). The IFOF appears to have lost a large portion of its superior-frontal lobe connections as a result of the resection, which is reflected in the damage rating increase (rating 1→2) and noted in the resected tissue missing from the post-surgery IFOF model (orange arrow). The Arcuate, however, did not change in subjective rating pre- vs. post-surgery (rating 2→2); the fascicle model was partially destroyed pre-surgery and futher destroyed post-surgery, but the rating system was not resolute enough to reflect that distinction. This case shows the advantage of having a continuous representation of damage, as opposed to a discrete one; the Arcuate is subjectively codified as unchanged, but the difference pipeline connectivity shows the specific connection that was threatened by the surgery.

**Figure 4:**
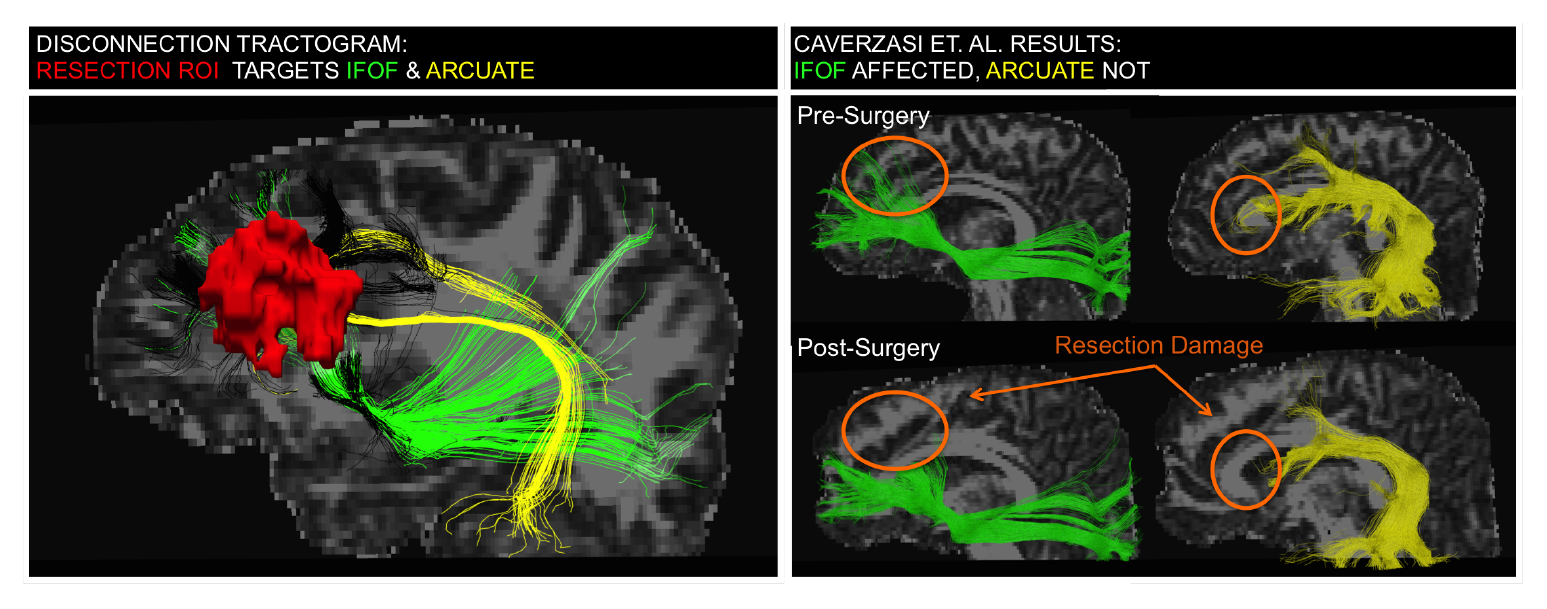
Continuous Damage Representation. From the connectivity of the ASAP ROI (red), portions of the left Arcuate Fascicle (yellow) and left Inferior Fronto-Occipital Fascicle (green) models were segmented. This agrees with the manual reconstructions of the pre- and post-surgery Arcuate and IFOF (RIGHT) models. The IFOF appears to have lost a big chunk to the resection (damage rating 1→2). The Arcuate, however, did not change in subjective rating pre- vs. post-surgery (rating 2→2); the fascicle model was partially destroyed pre-surgery and futher destroyed post-surgery, but the rating system was not resolute enough to reflect that distinction. Manual Results: IFOF (1→2), Uncinate (0→2), SLFIII (2→3, Arcuate (2→2), SLFII (1→1)

Another case that demonstrates the advantage of evaluating white matter disconnection on a continuous scale is shown in Figure 5 and Supplementary Figure 4M. According to the disconnection tractogram, the region immediately adjacent to the tumor is connected to the temporal lobe, frontal lobe, and spinal cord. However, these streamlines do not meet the criteria for inclusion in the any anatomically-constrained fascicle models because both ends do not reach the cortex. These bundles follow trajectories characteristic of the Arcuate Fascicle (yellow), the IFOF (green), and a bundle connected to the spinal cord (blue). The pre-surgical model of the Arcuate does not include the component wrapped around the tumor because it does not reach the cortex, so the pre- vs. post-surgical models were rated as unchanged.

**Figure 5:**
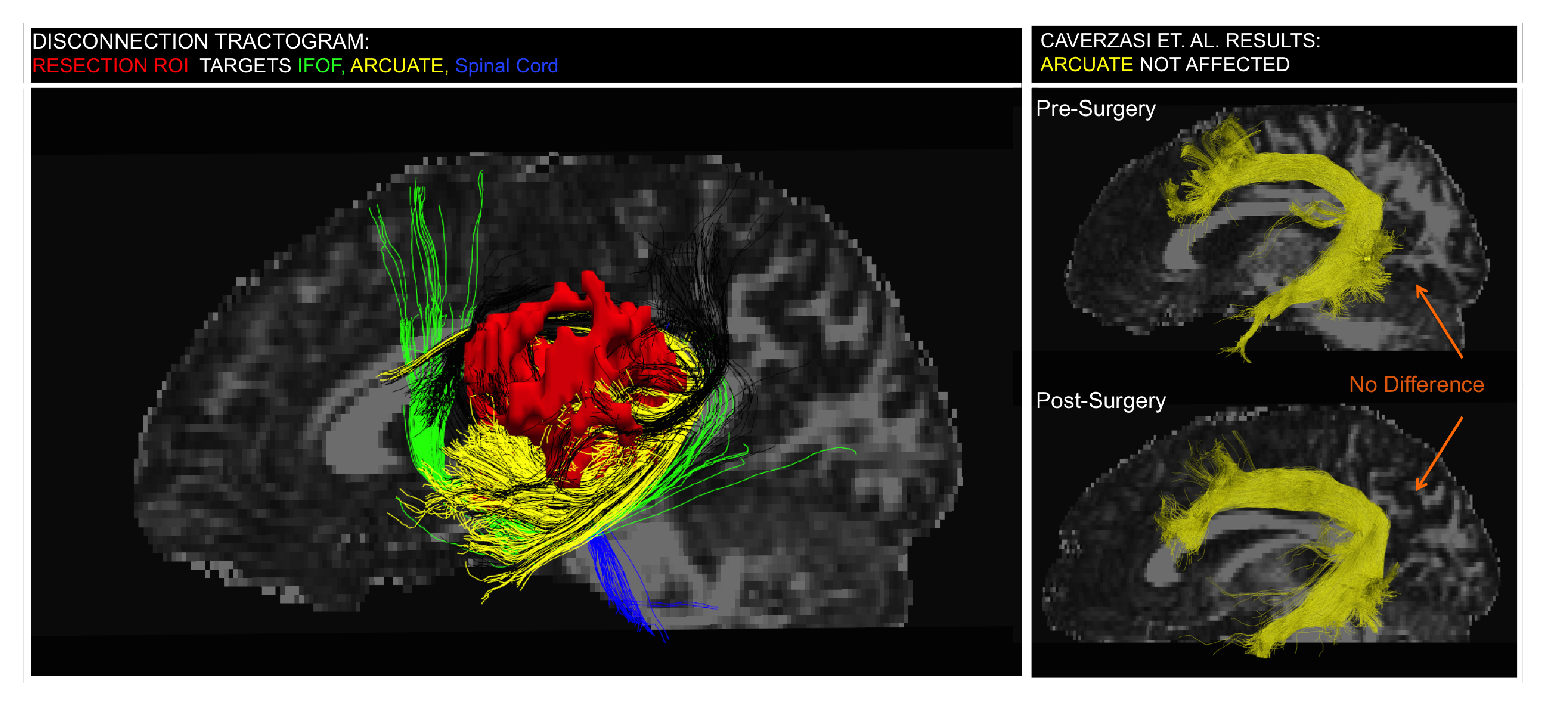
Wrapping Around the Tumor: Artifact or Important? The region immediately adjacent to the tumor contains tracks that do not meet the criteria for inclusion in the any Anatomically-Constrained fascicle models because both ends do not reach the cortex. However, the bundles follow trajectories characteristic of the Arcuate Fascicle (yellow), the IFOF (green), and a bundle connected to the spinal cord (blue). The pre-surgical model of the Arcuate does not include the component wrapped around the tumor because it does not reach the cortex, so the pre- vs. post-surgical models were rated as unchanged.

### 3.3 Performance Despite Distortion and Artifacts

The difference pipeline performs well, in spite of interference from artifacts or tissue distortion. The patient shown in Figure 6 had a particularly septated heterogeneous lesion with cystic, nodular and hemorrhagic components, which was further complicated by post-surgical hemorrhage exerting pressure on the tissue. A mass effect is apparent in the pre- and post-surgical images, but the pipeline still resulted in a reasonable output similar to the manual assessment by neuroradiologists. The output showed a sub-bundle of the Arcuate (yellow) as the likely white matter disconnection occurring as a result of the resection. The manual approach rated the only changes as Arcuate Fascicle and SLF II increasing in damage from 1→2. One point to note, in a case like this, performing tractography on the post-surgical dataset may be very difficult because of challenges associated with performing tractography post-surgery (e.g. edema, blood products, metallic implants, etc.). The cavity is not adjacent to the SLF II, so the damage rating to the SLF II structure is questionable; it is possible that the manual reconstruction was missing components due to post-surgery tracking difficulty, as opposed to the tissue being resected. The difference pipeline produces disconnection tractograms using pre-surgical data; the post-surgical data is only used to identify cavity, so surgery-related edema or blood products has less effect on the results.

**Figure 6:**
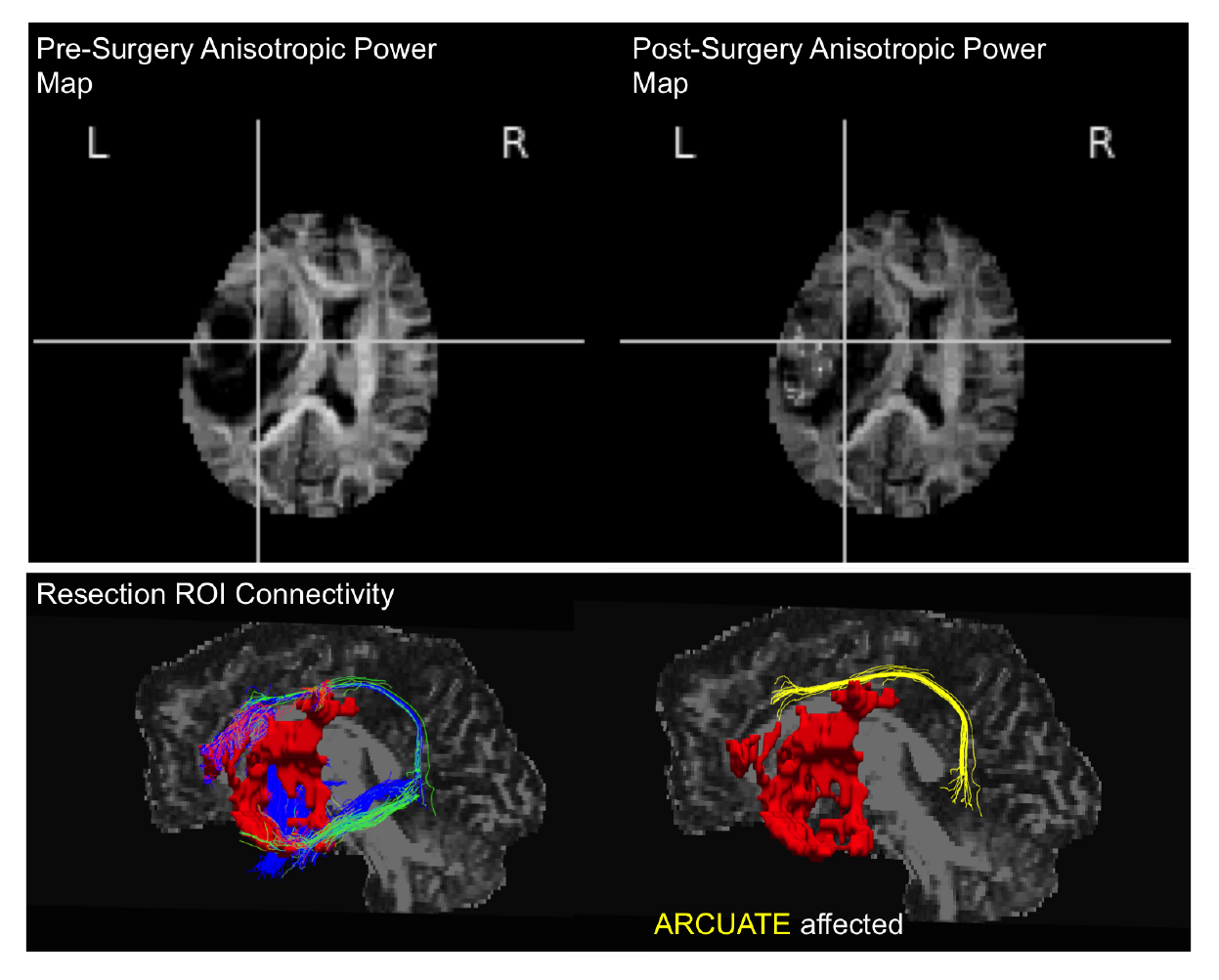
Performance with Severe Tissue Distortion. In this case, severe mass effect was observed due to the aggressive growth of the tumor and pressure from hemorrhaging. This pipeline was still able to estimate an ASAP ROI and produce a prediction that the Arcuate Fascicle was disconnected. The manual rating also cited the SLFII as being damaged, but the underlying white matter is intact so this discrepancy could be explained by post-surgical tracking difficulty. Manual Results: Arcuate (1→2), SLFII (1→2), SLFIII (2→2), SLF-tp (3→3), ILF (3→3), MdLF (3→3), IFOF (3→3), Uncinate (3→3), CST face/hand/foot (1→1)

## 4 Discussion

The ASAP-Tractogram approach, presented here, meets the need for an objective, automatic pipeline robust to pathology. The objective and automatic aspects have vast implications on scalability of studies and translation across institutions, but the pipeline approach, itself, has unique advantages for studies of any size. Sub-bundle resolution is important in executing specific studies relating white matter connectivity to functional deficits; the classification of a subcortical pathway may include a variety of connections (Fernández-Miranda et al. 2014) and there may not be a consensus on the exact components of an anatomically-defined fascicle model (Heide et al. 2013). Also, there is evidence that a bundle’s robustness to damage is not uniform (Herbet et al. 2016).

The need for technical development of tractography methods is widely acknowledged (Duffau 2014), particularly to eliminate manual intervention, which creates a danger for excluding displaced fibers that represent genuinely displaced white matter bundles (Feigl et al. 2014). We found this concern to be justified in this study, as well. As an example, the case presented in Figure 5 had subjective ratings of fascicle damage that did not explain the permanent new language deficit the patient experienced. An analysis of the difference pipeline results suggested that a fragment of the model of the Arcuate Fascicle, a white matter structure associated with language deficits (Bernal and Ardila 2009), wrapped around the tumor and had potentially been disrupted by the surgery. While this wrapping phenomenon is not well-characterized, it is reasonable to consider that information can be gleaned from tractography models engaged in wrapping merely because of their proximity to the tumor, which puts the fascicles they represent at risk of being damaged by the surgical resection. In the manual fascicle modeling of this case, streamlines that wrapped around the tumor region were removed by the manual operator because they did not match the anatomical constraints of the Arcuate Fascicle, so the potential disruption was not reflected by the post-operative rating scale because the at-risk bundle was excluded from the pre-operative model (Supplementary Figure 4M). In the case shown in Figure 5, a distorted fascicle model of a bundle indicating connectivity to the spinal cord was wrapped around the tumor, but did not reach the cortex so the bundle classification is inconclusive. The patient had a short-term post-surgical motor deficit, so information about connectivity to the spinal cord may be relevant to this case. It may be that the white matter directly adjacent to the tumor region helps to explain the discrepancy between tissue damage and functional deficits, but tractography models in that tissue are highly uncertain due to the presence of disease processes.

This pipeline approach is also robust to pathology, provided that distortions are similar between time-points. The approach only attempts to precisely register images longitudinally; there is no registration to MNI. Cases, such as Figure 6, may be excluded from VLSM studies due to the extreme level of tissue distortion (Herbet et al. 2016) but the distortion is similar enough between pre- and post-surgery to analyze the case longitudinally. The resulting tractography model generated on the single-subject level can be compared in a cohort study without ever registering the distorted tissue, itself, to a common space. This pipeline models the connectivity of the resection tissue in the patient-specific space, which confers higher confidence in the identity of the white matter structure threatened by the surgery than comparison to healthy controls (Almairac et al. 2014).

### 4.1 Limitations of the ASAP-Tractogram Approach

While there are many advantages conferred by this automatic image processing pipeline approach, a number of limitations must be carefully considered throughout the implementation and use of this workflow. Resection studies are very difficult, from an engineering standpoint, because the mechanics of tissues are difficult to predict (Dixit and Liu 2016; Tse et al. 2013), especially when a heterogeneous pathology is involved (Garlapati et al. 2014). There is a significant concern for how well the tissues can be matched from pre- vs. post-surgery imaging and at what point correcting for the tissue shifts that come with focal lesion resections (the previous tumor growth exerting local pressure that subsequently relaxes back into the resection cavity after removal, which eventually disappears), gravity causing tissue to “cave in” to larger resections, blood causing both imaging artifacts and, in some cases, exerting pressure on the brain, implants causing imaging artifacts, air causing both tissue displacement and imaging artifacts, pressure/CSF changes, swelling of tissue, to name a few. Setting parameters for longitudinal registration between peri-surgical time points involves a tradeoff: the nonlinear registration algorithm must reflect genuine tissue shift associated with the surgery, but it cannot be allowed so much flexibility that it “corrects” the surgical cavity.

Another consideration when deciding on an experimental approach is when and how to collect the HARDI datasets. There is a tradeoff between some of the challenges present immediately postop, and the risk of tissue reorganization or further disease processes occurring at a later time-point, after immediate post-operative challenges have resolved. One of the aspects of post-surgical images that makes registration to pre-surgical images difficult in this particular cohort is the susceptibility artifacts caused by blood, air, and implants being imaged using an Echo-Planar sequence. The HARDI datasets from this study were not collected with any components necessary for susceptibility correction at the time of acquisition. Susceptibility correction can be trivial, provided either a field-map (Jezzard and Balaban 1995) or two B0 images with opposite phase-encoding gradients are collected at the time of acquisition (Andersson, Skare, and Ashburner 2003). Applying this pipeline to susceptibility-corrected images should improve performance, as that removes one element of mismatch pre vs. post-surgery. The authors strongly recommend that, if possible, susceptibility correction is considered in sequence planning.

This connectivity-focused approach may need to be complemented by a VLSM study to better explain the deficits in a cohort. Some studies have shown that it is important to consider the location of damage along the length of the track (Herbet et al. 2016). Also, only damage associated with spatially coherent decreases in AP will be identified by this approach. It may be possible to optimize this pipeline for other types of injury. For example, known changes in mean diffusivity (MD) associated with ischemic injury could be leveraged to segment potential regions of infarct.

### 4.2 Artifacts and Errors

One important limitation of any new image-processing pipeline is that the output must be scrutinized critically for new artifacts and errors. In this cohort, the heterogeneity of the parameter changes around the surgical cavity presented some unexpected results. In most cases, the pattern of AP decreasing and MD increasing pre- to post-surgery was reversed in some regions around the surgical site. AP is extremely hyperintense in these regions postsurgery (greater than the intensity of the corpus-callosum, which should have some of the highest AP in the whole brain) (Example: Figure 6). Based on comparison by a neuroradiologist to CT images, there were two explanations for this phenomenon: pneumocephalus and blood products. When air is present inside the skull (pneumocephalus), MD decreases because CSF/tissue is displaced by air, so the AP presents as artifactually hyperintense post-surgery resulting in a net increase in the AP difference parameter. This parameter reversal was apparent in several cases, as can be seen in Supplementary Figure 5. AP is increasing and MD is decreasing, because AP appears to be artifactually hyperintense in the presence of blood. The decrease in MD is logical because blood is viscous, restricting diffusion. Anisotropic power is a relatively new measurement, so it has yet to be characterized. This artifact in AP hyperintensity may be due to the dephasing of signal in the B0 image due to inhomogeneity of fields characteristic of blood because of its high paramagnetic properties and air/tissue interface creates a susceptibility artifact.

The most common mode of QC failure was due to the presence of false-positive resection ROI’s. The cause of this tended to be an error in the automatic segmentation of the AP change due to excessive distortion of ventricles. Blood products, pneumocephalus, and other distortions related to the surgery can cause the ventricles to shift, in relation to the preoperative image. In these cases, the peri-ventricular white matter is mistaken for a resection ROI because parts of the highly anisotropic corpus callosum overlay the shifted ventricle in the post-operative image. This phenomenon caused two patients in the cohort to fail QC, but was easy to identify and correct.

In one case, the difference pipeline missed some bundles because they were not represented in the pre-surgery streamline dataset used for targeting the resection ROI.

This is due to insufficient seeding density; these streamlines were represented in the (Caverzasi et al. 2016) study because the seeding density was much higher.

Many concerns about using this ASAP pipeline approach can be addressed by rigorous implementation standards: a pilot study should always be performed with the results from a variety of extreme patient conditions evaluated by neuroradiologists, and a strict QC protocol employed. Several features of diffusion parameters lend themselves to serving as indicators for QC.

### 4.3 Potential to Fully Automate Disconnection Studies

This pipeline automates the generation of a disconnection tractogram associated with focal injury, but an expert still had to classify the ASAP-Tractogram components by recognizing the trajectory of the pathway. Automatic classification algorithms are being advanced continuously; some have even been demonstrated as robust to some pathology (Yeatman et al. 2012; O’Donnell et al. 2017). The combination of this ASAP-Tractogram approach with automatic classification algorithms would enable fully automatic studies. This could facilitate Pathway Lesion Symptom Mapping studies characterizing the risk of a particular deficit with the confidence that comes from very large studies. These high-confidence studies are necessary to empower clinicians to ask the question “What is the possible morbidity to this unique patient given the location, size, and extent of this infiltrative tumor?” so that they can better counsel their patients and plan interventions to minimize complications and maximize the extent of resection, which will impact patient quality and quantity of life.

This pipeline could also be applied to other types of longitudinal white matter damage. Neurosurgical intervention in pathologies involving less tissue distortion should work at least as well as this tumor demonstration (the pre-surgical brains in non-tumor epilepsy patients should be more similar to controls than, for example, a grade IV tumor). Pathologies like stroke and Multiple Sclerosis may require tuning of parameters but, in theory, should be a good application for this pipeline because the deformations between time points tend to be smaller than those associated with tumor resection.

### 4.4 Potential to Impact Clinical Decisions

A shift in the framework of thinking about the brain has been progressing from a topographical approach to one more focused on pathways (“hodotopical”)(Catani 2005). This is reflected in efforts by the field of neurosurgery to move beyond preservation of discrete “eloquent regions” toward a holistic approach integrating information from many anatomical/functional systems (Duffau 2015). Tractography methods are being pushed into the neurosurgical navigation systems because they provide in-vivo models of patient-specific white matter structures, essential to understanding this new paradigm. There is growing evidence that the potential of functional compensation for white matter damage is limited (Ius et al. 2011; He et al. 2007; Genova et al. 2014; Cristofori et al. 2015) and that post-surgical plasticity relies on long-range white matter connections being spared by surgery (Duffau 2014). There is evidence that damage to a functional system can be compensated for by adjacent regions (Benzagmout, Gatignol, and Duffau 2007) and that long-range connections may help cortical reorganization (Papagno et al. 2011). All of these questions are important to answer for both basic neuroscience and applied clinical researchers, but the studies required to develop confidence in the answers cannot be done with today’s technology. To further advance studies relating focal white matter damage to deficits and translatability of results across institutions, methods that are objective, automatic, and compatible with brains distorted by pathology are necessary.

## 5 Conclusions

The proposed ASAP-tractogram approach may provide advantages over current methods for both accuracy and efficiency when confronted with pathology that significantly distorts the normal anatomy (as is often the case for brain tumors). The extent of variability due to genetics, learning over a lifetime, varying functional states, pathological processes, the brain’s response to pathology (Duffau 2013), and uncertainty in models of structural/functional systems necessitates studies with considerably larger sample sizes and more specific characterization of disrupted white matter structures than are the standard with current technologies. Our work lays the groundwork for high-resolution, larger scale, sufficiently powered analyses by meeting the need for an automatic approach to objectively quantify white matter disconnection.

## Acknowledgements

The authors would like to thank Valentina Panara, Shawn Hervey-Jumper, Caroline Racine, Francesco Sanvito, Simone Sacco, Ariel Rokem, Antonella Castellano, Robert Knight, Pratik Mukherjee, Vanitha Sankaranarayanan, and all of our research subjects.

## 6 Appendices

**Supplementary Figure 1:**
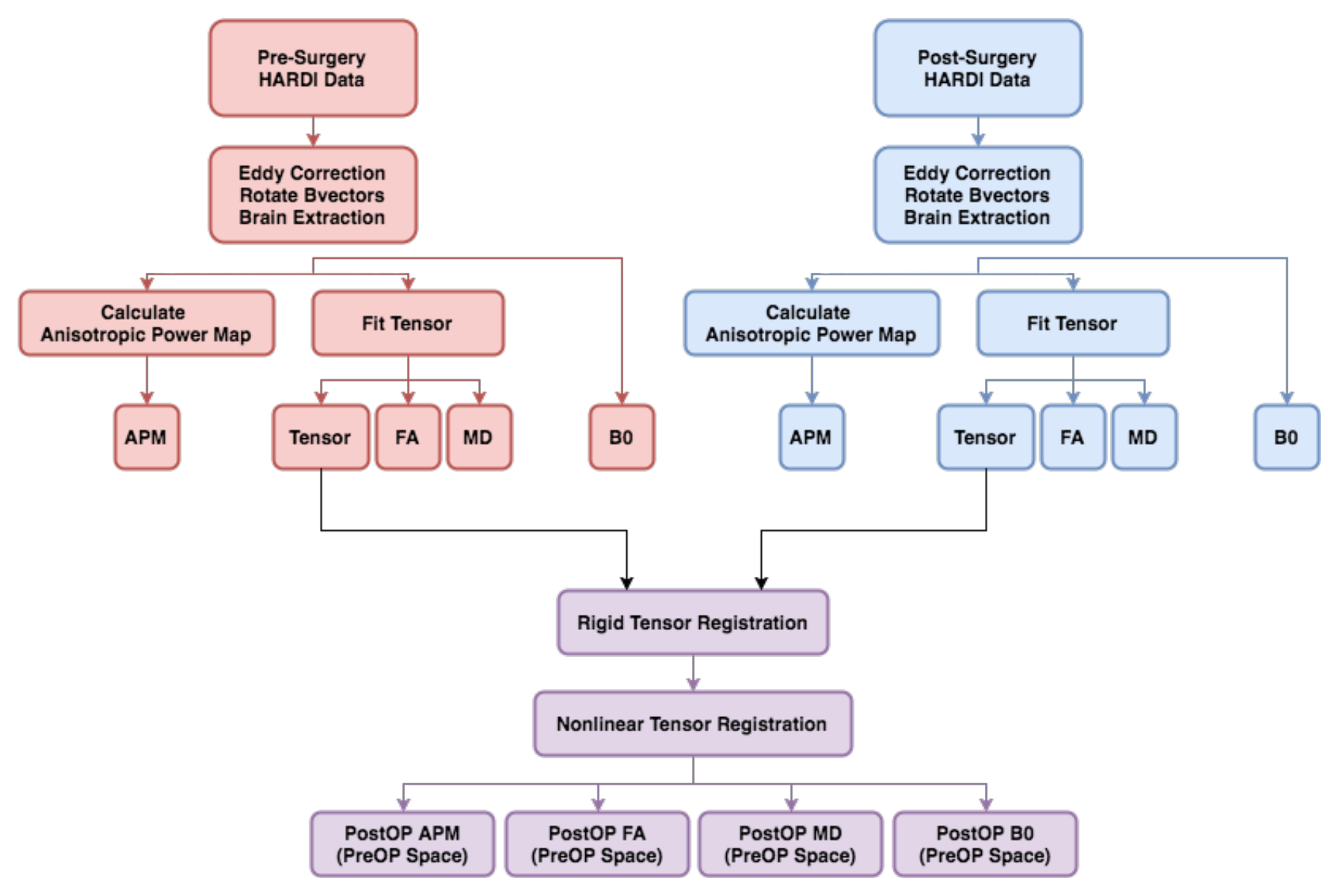
A flowchart of the preprocessing steps executed to produce postsurgery diffusion metric images aligned to the pre-surgery images. (APM = Anisotropic Power Map, FA = Fractional Anisotropy, MD = Mean Diffusivity, B0 = minimally diffusion-weighted image)

**Supplementary Figure 2:**
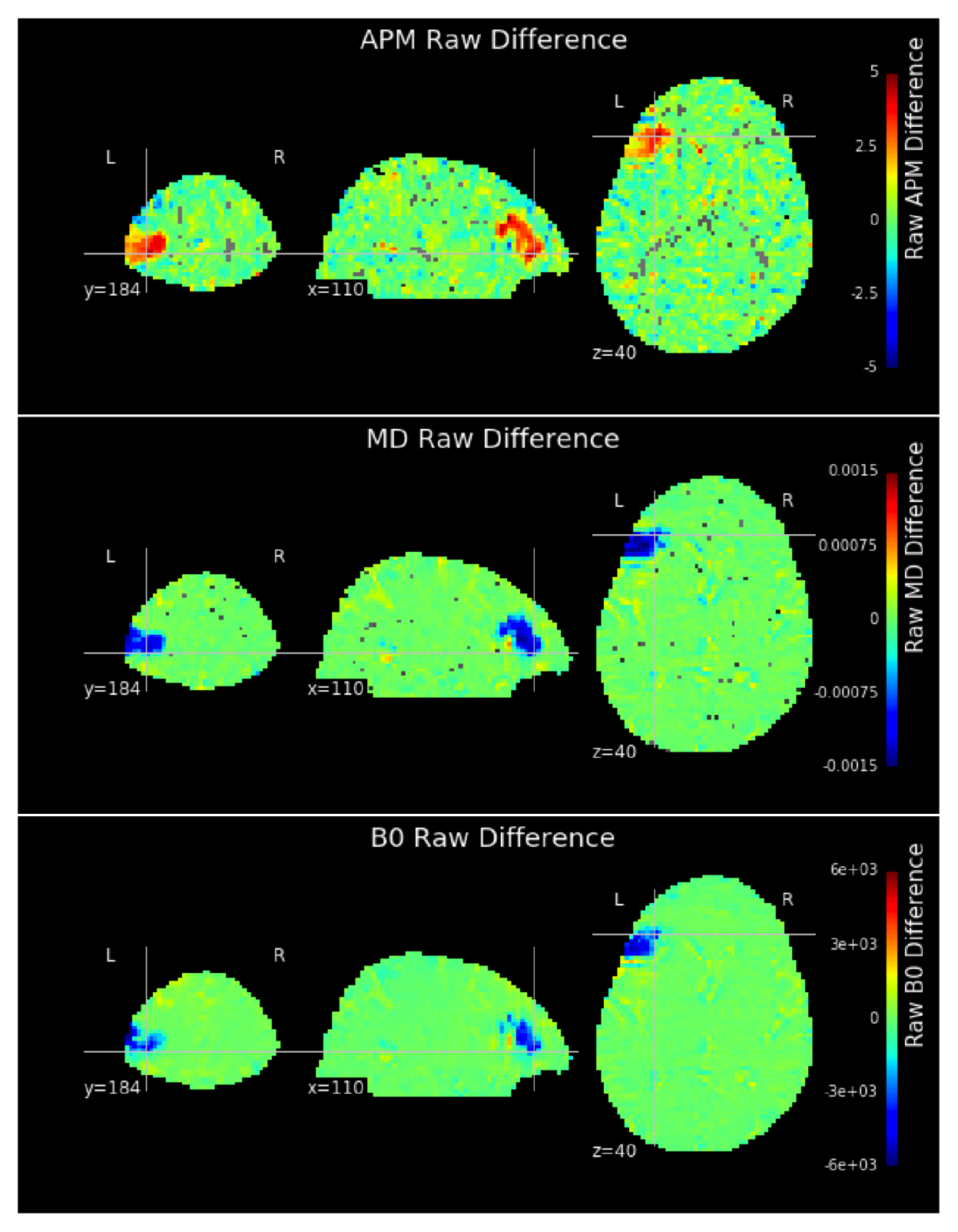
Difference maps were created by subtracting the aligned post-surgical diffusion metric image from the pre-surgical. The resection cavity is shown as positive in the anisotropic power map (APM) because anisotropic tissue was removed, and negative in the mean diffusivity (MD) and B0 maps because tissue was replaced with CSF, which has higher diffusivity and T2-contrast intensity.

**Supplementary Figure 3:**
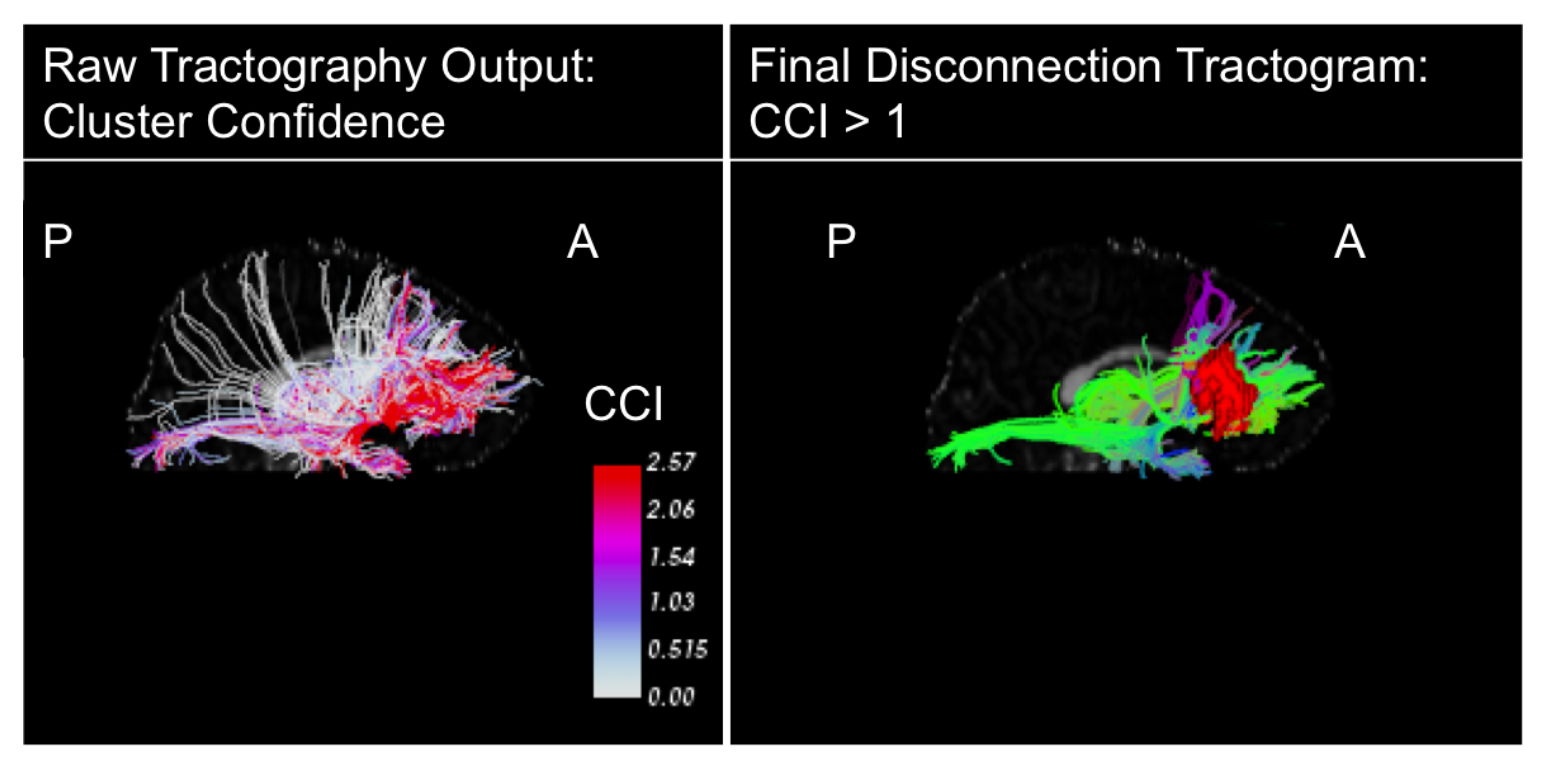
The streamline output was filtered objectively using the Cluster Confidence Index (CCI), to exclude low-confidence streamlines (left panel). Any streamlines with CCI <1 or length <40mm were excluded from analysis, resulting in the final disconnection tractogram (right panel), shown with the ASAP ROI.

**Supplementary Figure 4:**
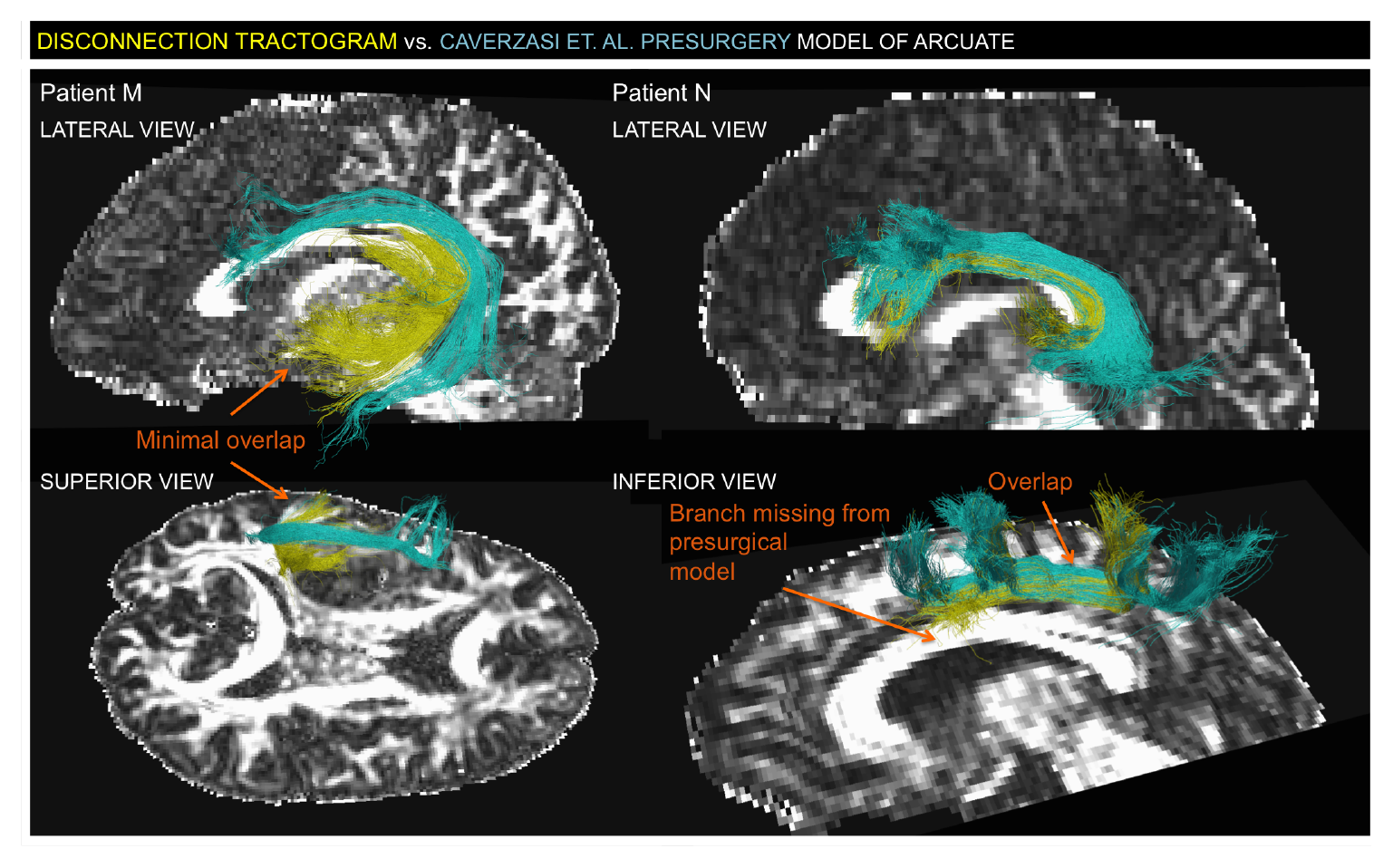
Disconnection Tractogram vs. ACT Presurgical Model of Arcuate Fascicle in Patients with Postsurgical Language Deficits. The presurgical model of the Arcuate Fascicle (Blue) in Patient M (LEFT) is almost entirely distinct from the disconnection tractogram (yellow), which shows a fragment of an Arcuate Fascicle model wrapped around the tumor region. The presurgical model of the Arcuate Fascicle (Blue) in Patient N (RIGHT) largely overlaps with the presurgical model, evidenced by the interdigitation of the blue and yellow streamlines.

**Supplementary Figure 5:**
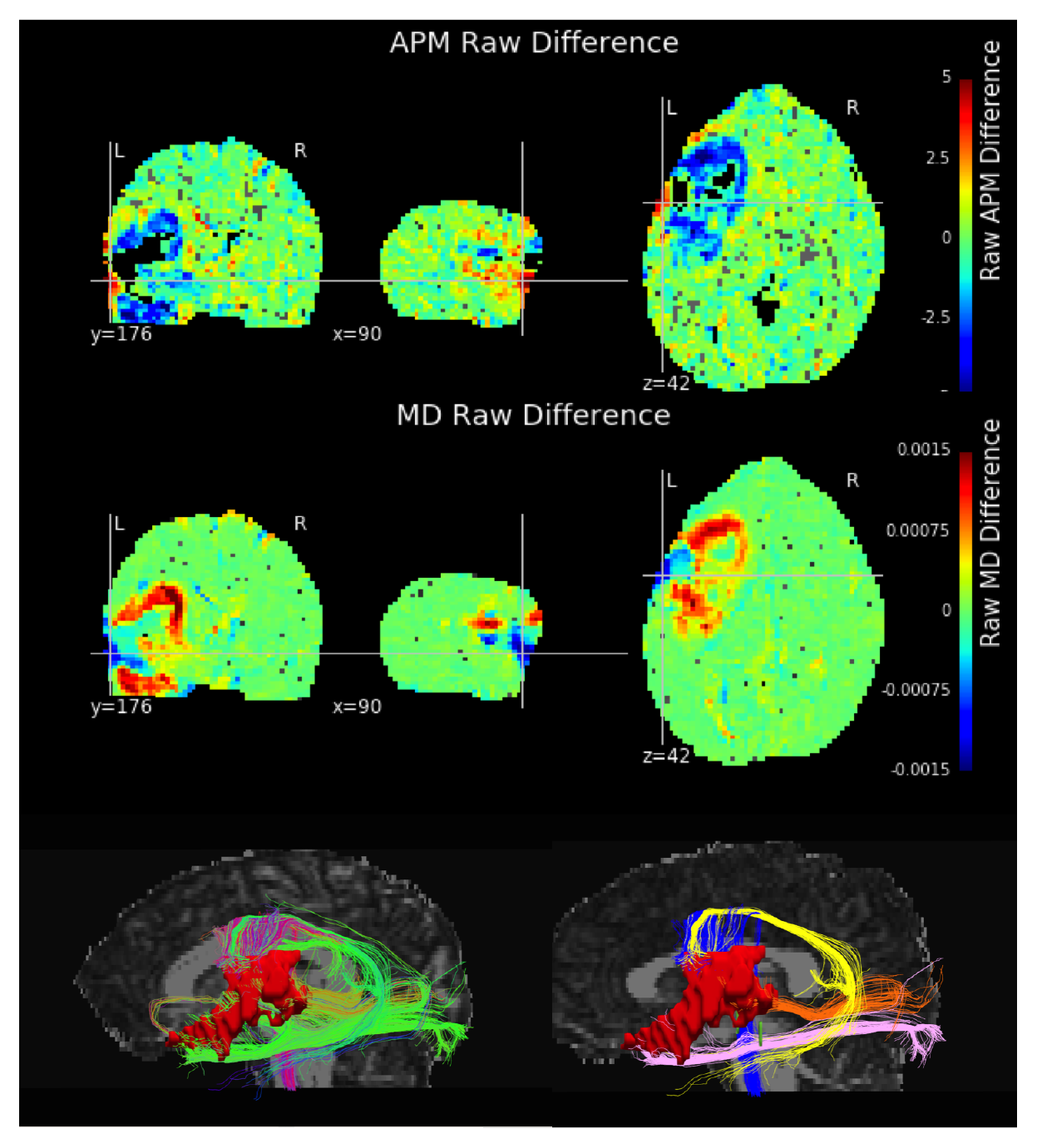
Blood Products and Pneumocephalus. This patient had pneumocephalus and blood products which appears to manifest as inverted contrast on the anisotropic power map (APM) and mean diffusivity (MD). A) Blood and pneumocephalus are apparently associated with hyperintense AP, resulting in an artifactual increase in AP postsurgery. B) Mean diffusivity decreases due to viscous blood, air, and ischemia. C) The difference pipeline still succeeds in isolating the tissue that decreased in AP. This difference map suggsts that the Arcuate (yellow), Medial Longitudinal Fascicle (MdLF=orange), the Inferior Longitudinal Fascicle (ILF), and a bundle connected to the spinal cord (blue) were at risk of being disrupted by the surgical resection. Manual Results: Arcuate (1→2), MdLF (1→3), ILF (2→2), SLFII (2→2), SLFIII (3→3), IFOF (3→3), Uncinate (3→3), CST face/hand (1→1)

## References

Berti, Anna, Francesca Garbarini, and Marco Neppi-Modona. 2015. “Disorders of Higher Cortical Function”. In Neurobiology of Brain Disorders, 525–41. Elsevier BV. doi:10.1016/b978-0-12-398270-4.00032-x.

Geschwind, Norman. 1965. “Disconnexion Syndromes in Animals and Man”. Brain 88 (2). Oxford University Press (OUP): 237–94. doi:10.1093/brain/88.2.237.

Bates, Elizabeth, Stephen M. Wilson, Ayse Pinar Saygin, Frederic Dick, Martin I. Sereno, Robert T. Knight, and Nina F. Dronkers. 2003. “Voxel-Based LesionSymptom Mapping”. Nature Neuroscience, April. Springer Nature. doi:10.1038/nn1050.

Geschwind, Norman. 1965. “Disconnexion Syndromes in Animals and Man”. Brain 88 (3). Oxford University Press (OUP): 585–644. doi:10.1093/brain/88.3.585.

Basser, P.J., J. Mattiello, and D. LeBihan. 1994. “MR Diffusion Tensor Spectroscopy and Imaging”. Biophysical Journal 66 (1). Elsevier BV: 259–67. doi:10.1016/s0006-3495(94)80775-1.

Le Bihan, Denis, Jean-François Mangin, Cyril Poupon, Chris A Clark, Sabina Pappata, Nicolas Molko, and Hughes Chabriat. 2001. “Diffusion Tensor Imaging: Concepts and Applications”. Journal of Magnetic Resonance Imaging 13 (4). Wiley Online Library: 534–46.

Pierpaoli, C, P Jezzard, P J Basser, A Barnett, and G Di Chiro. 1996. “Diffusion Tensor MR Imaging of the Human Brain.”. Radiology 201 (3). Radiological Society of North America (RSNA): 637–48. doi:10.1148/radiology.201.3.8939209.

Pajevic, Sinisa, and Carlo Pierpaoli. 1999. “Color Schemes to Represent the Orientation of Anisotropic Tissues from Diffusion Tensor Data: Application to White Matter Fiber Tract Mapping in the Human Brain”. Magnetic Resonance in Medicine 42 (3). Wiley-Blackwell: 526–40. doi:10.1002/(sici)1522-2594(199909)42:3<526::aid-mrm15>3.3.co;2-a.

Tuch, David S. 2004. “Q-Ball Imaging”. Magnetic Resonance in Medicine 52 (6). Wiley-Blackwell: 1358–72. doi:10.1002/mrm.20279.

Tournier, J.-Donald, Fernando Calamante, David G. Gadian, and Alan Connelly. 2004. “Direct Estimation of the Fiber Orientation Density Function from Diffusion-Weighted MRI Data Using Spherical Deconvolution”. NeuroImage 23 (3). Elsevier BV: 1176–85. doi:10.1016/j.neuroimage.2004.07.037.

Jensen, Jens H., Joseph A. Helpern, Anita Ramani, Hanzhang Lu, and Kyle Kaczynski. 2005. “Diffusional Kurtosis Imaging: The Quantification of Non-Gaussian Water Diffusion by Means of Magnetic Resonance Imaging”. Magnetic Resonance in Medicine 53 (6). Wiley-Blackwell: 1432–40. doi:10.1002/mrm.20508.

Assaf, Yaniv, and Peter J. Basser. 2005. “Composite Hindered and Restricted Model of Diffusion (CHARMED) MR Imaging of the Human Brain”. NeuroImage 27 (1). Elsevier BV: 48–58. doi:10.1016/j.neuroimage.2005.03.042.

Jeurissen, Ben, Jacques-Donald Tournier, Thijs Dhollander, Alan Connelly, and Jan Sijbers. 2014. “Multi-Tissue Constrained Spherical Deconvolution for Improved Analysis of MultiShell Diffusion MRI Data”. NeuroImage 103 (December). Elsevier BV: 411–26. doi:10.1016/j.neuroimage.2014.07.061.

Tuch, David S., Timothy G. Reese, Mette R. Wiegell, Nikos Makris, John W. Belliveau, and Van J. Wedeen. 2002. “High Angular Resolution Diffusion Imaging Reveals Intravoxel White Matter Fiber Heterogeneity”. Magnetic Resonance in Medicine 48 (4). Wiley-Blackwell: 577–82. doi:10.1002/mrm.10268.

Hess, Christopher P., Pratik Mukherjee, Eric T. Han, Duan Xu, and Daniel B. Vigneron. 2006. “Q-Ball Reconstruction of Multimodal Fiber Orientations Using the Spherical Harmonic Basis”. Magnetic Resonance in Medicine 56 (1). Wiley-Blackwell: 104–17. doi:10.1002/mrm.20931.

Kumar, Pradeep, Prachi Kathuria, Pallavi Nair, and Kameshwar Prasad. 2016. “Prediction of Upper Limb Motor Recovery after Subacute Ischemic Stroke Using Diffusion Tensor Imaging: a Systematic Review and Meta-Analysis”. Journal of Stroke 18 (1). Korean Stroke Society: 50.

Kumar, Pradeep, Arun Kumar Yadav, Shubham Misra, Amit Kumar, Kamalesh Chakravarty, and Kameshwar Prasad. 2016. “Prediction of Upper Extremity Motor Recovery after Subacute Intracerebral Hemorrhage through Diffusion Tensor Imaging: a Systematic Review and Meta-Analysis”. Neuroradiology 58 (10). Springer: 1043–50.

Yogarajah, M., N. K. Focke, S. Bonelli, M. Cercignani, J. Acheson, G. J. M. Parker, D. C. Alexander, et al. 2009. “Defining Meyers Loop-Temporal Lobe Resections Visual Field Deficits and Diffusion Tensor Tractography”. Brain 132 (6). Oxford University Press (OUP): 1656–68. doi:10.1093/brain/awp114.

Chen, Xiaolei, Daniel Weigel, Oliver Ganslandt, Michael Buchfelder, and Christopher Nimsky. 2009. “Prediction of Visual Field Deficits by Diffusion Tensor Imaging in Temporal Lobe Epilepsy Surgery”. NeuroImage 45 (2). Elsevier BV: 286–97. doi:10.1016/j.neuroimage.2008.11.038.

Glenn, Orit A., Roland G. Henry, Jeffrey I. Berman, Patrick C. Chang, Steven P. Miller, Daniel B. Vigneron, and A. James Barkovich. 2003. “DTI-Based Three-Dimensional Tractography Detects Differences in the Pyramidal Tracts of Infants and Children with Congenital Hemiparesis”. Journal of Magnetic Resonance Imaging 18 (6). Wiley-Blackwell: 641–48. doi:10.1002/jmri.10420.

Mandelli, M. L., E. Caverzasi, R. J. Binney, M. L. Henry, I. Lobach, N. Block, B. Amirbekian, et al. 2014. “Frontal White Matter Tracts Sustaining Speech Production in Primary Progressive Aphasia”. Journal of Neuroscience 34 (29). Society for Neuroscience: 9754–67. doi:10.1523/jneurosci.3464-13.2014.

Jang, Sung Ho. 2013. “Diffusion Tensor Imaging Studies on Arcuate Fasciculus in Stroke Patients: A Review”. Frontiers in Human Neuroscience 7. Frontiers Media SA. doi:10.3389/fnhum.2013.00749.

Mori, Susumu, and Peter van Zijl. 2002. “Fiber Tracking: Principles and Strategies-a Technical Review”. NMR in Biomedicine 15 (7–8). Wiley Online Library: 468–80.

Caverzasi, Eduardo, Shawn L. Hervey-Jumper, Kesshi M. Jordan, Iryna V. Lobach, Jing Li, Valentina Panara, Caroline A. Racine, et al. 2016. “Identifying Preoperative Language Tracts and Predicting Postoperative Functional Recovery Using HARDI q-Ball Fiber Tractography in Patients with Gliomas”. Journal of Neurosurgery 125 (1). Journal of Neurosurgery Publishing Group (JNSPG): 33–45. doi:10.3171/2015.6.jns142203.

Duffau, Hugues. 2008. “The Anatomo-Functional Connectivity of Language Revisited”. Neuropsychologia 46 (4). Elsevier BV: 927–34. doi:10.1016/j.neuropsychologia.2007.10.025.

Kim, S. H., and S. H. Jang. 2012. “Prediction of Aphasia Outcome Using Diffusion Tensor Tractography for Arcuate Fasciculus in Stroke”. American Journal of Neuroradiology 34 (4). American Society of Neuroradiology (ASNR): 785–90. doi:10.3174/ajnr.a3259.

Wu, Jin-Song, Liang-Fu Zhou, Wei-Jun Tang, Ying Mao, Jin Hu, Yan-Yan Song, Xun-Ning Hong, and Gu-Hong Du. 2007. “CLINICAL EVALUATION AND FOLLOW-UP OUTCOME OF DIFFUSION TENSOR IMAGING-BASED FUNCTIONAL NEURONAVIGATION”. Neurosurgery 61 (5). Oxford University Press (OUP): 935–49. doi:10.1227/01.neu.0000303189.80049.ab.

Leclercq, Delphine, Hugues Duffau, Christine Delmaire, Laurent Capelle, Peggy Gatignol, Mathieu Ducros, Jacques Chiras, and Stéphane Lehéricy. 2010. “Comparison of Diffusion Tensor Imaging Tractography of Language Tracts and Intraoperative Subcortical Stimulations”. Journal of Neurosurgery 112 (3). Journal of Neurosurgery Publishing Group (JNSPG): 503–11. doi:10.3171/2009.8.jns09558.

Henry, Roland G, Jeffrey I Berman, Srikantan S Nagarajan, Pratik Mukherjee, and Mitchel S Berger. 2004. “Subcortical Pathways Serving Cortical Language Sites: Initial Experience with Diffusion Tensor Imaging Fiber Tracking Combined with Intraoperative Language Mapping”. NeuroImage 21 (2). Elsevier BV: 616–22. doi:10.1016/j.neuroimage.2003.09.047.

Lehéricy, Stéphane, Hugues Duffau, Pierre-François Van de Moortele, and Christine Delmaire. 2007. “Validity of Presurgical Functional Localization”. In Clinical Functional MRI, 167–87. Springer Nature. doi:10.1007/978-3-540-49976-3_7.

Lehéricy, Stéphane, Delphine Leclercq, Hugues Duffau, Pierre-François Van de Moortele, and Christine Delmaire. 2014. “Presurgical Functional Localization Possibilities Limitations, and Validity”. In Clinical Functional MRI, 247–67. Springer Nature. doi:10.1007/978-3-662-45123-6_9.

Berman, Jeffrey I., Mitchel S. Berger, Sungwon Chung, Srikantan S. Nagarajan, and Roland G. Henry. 2007. “Accuracy of Diffusion Tensor Magnetic Resonance Imaging Tractography Assessed Using Intraoperative Subcortical Stimulation Mapping and Magnetic Source Imaging”. Journal of Neurosurgery 107 (3). Journal of Neurosurgery Publishing Group (JNSPG): 488–94. doi:10.3171/jns-07/09/0488.

Bucci, Monica, Maria Luisa Mandelli, Jeffrey I. Berman, Bagrat Amirbekian, Christopher Nguyen, Mitchel S. Berger, and Roland G. Henry. 2013. “Quantifying Diffusion MRI Tractography of the Corticospinal Tract in Brain Tumors with Deterministic and Probabilistic Methods”. NeuroImage: Clinical 3. Elsevier BV: 361–68. doi:10.1016/j.nicl.2013.08.008.

Jordan, Kesshi, Bagrat Amirbekian, Anisha Keshavan, Roland Henry. “Cluster Confidence Index: A Streamline-wise Pathway Reproducibility Metric for Diffusion-Weighted MRI Tractography”. Journal of Neuroimaging (Accepted).

Mandelli, Maria Luisa, Mitchel S. Berger, Monica Bucci, Jeffrey I. Berman, Bagrat Amirbekian, and Roland G. Henry. 2014. “Quantifying Accuracy and Precision of Diffusion MR Tractography of the Corticospinal Tract in Brain Tumors”. Journal of Neurosurgery 121 (2). Journal of Neurosurgery Publishing Group (JNSPG): 349–58. doi:10.3171/2014.4.jns131160.

Berman, Jeffrey I. 2013. “Advanced Diffusion MR Tractography for Surgical Planning”. In Functional Brain Tumor Imaging, 183–94. Springer Nature. doi:10.1007/978-1-4419-5858-7_11.

Kinoshita, Manabu, Kei Yamada, Naoya Hashimoto, Amami Kato, Shuichi Izumoto, Takahito Baba, Motohiko Maruno, Tsunehiko Nishimura, and Toshiki Yoshimine. 2005. “Fiber-Tracking Does Not Accurately Estimate Size of Fiber Bundle in Pathological Condition: Initial Neurosurgical Experience Using Neuronavigation and Subcortical White Matter Stimulation”. NeuroImage 25 (2). Elsevier BV: 424–29. doi:10.1016/j.neuroimage.2004.07.076.

Duffau, Hugues. 2014. “The Dangers of Magnetic Resonance Imaging Diffusion Tensor Tractography in Brain Surgery”. World Neurosurgery 81 (1). Elsevier BV: 56–58. doi:10.1016/j.wneu.2013.01.116.

Chamberland, Maxime, Kevin Whittingstall, David Fortin, David Mathieu, and Maxime Descoteaux. 2014. “Real-Time Multi-Peak Tractography for Instantaneous Connectivity Display”. Frontiers in Neuroinformatics 8 (May). Frontiers Media SA. doi:10.3389/fninf.2014.00059.

Fillard, Pierre, Maxime Descoteaux, Alvina Goh, Sylvain Gouttard, Ben Jeurissen, James Malcolm, Alonso Ramirez-Manzanares, et al. 2011. “Quantitative Evaluation of 10 Tractography Algorithms on a Realistic Diffusion MR Phantom”. NeuroImage 56 (1). Elsevier BV: 220–34. doi:10.1016/j.neuroimage.2011.01.032.

Crettenand, S., S.D. Meredith, M.J. Hoptman, and R.B. Reilly. 2006. “Quantitative Analysis and Comparison of Diffusion Tensor Imaging Tractography Algorithms”. In IET Irish Signals and Systems Conference (ISSC 2006). Institution of Engineering and Technology (IET). doi:10.1049/cp:20060421.

Neher, Peter F., Maxime Descoteaux, Jean-Christophe Houde, Bram Stieltjes, and Klaus H. Maier-Hein. 2015. “Strengths and Weaknesses of State of the Art Fiber Tractography Pipelines A Comprehensive in-Vivo and Phantom Evaluation Study Using Tractometer”. Medical Image Analysis 26 (1). Elsevier BV: 287–305. doi:10.1016/j.media.2015.10.011.

Wakana, Setsu, Arvind Caprihan, Martina M. Panzenboeck, James H. Fallon, Michele Perry, Randy L. Gollub, Kegang Hua, et al. 2007. “Reproducibility of Quantitative Tractography Methods Applied to Cerebral White Matter”. NeuroImage 36 (3). Elsevier BV: 630–44. doi:10.1016/j.neuroimage.2007.02.049.

Golby, Alexandra J, Gordon Kindlmann, Isaiah Norton, Alexander Yarmarkovich, Steven Pieper, and Ron Kikinis. 2011. “Interactive Diffusion Tensor Tractography Visualization for Neurosurgical Planning”. Neurosurgery 68 (2). Oxford University Press (OUP): 496–505. doi:10.1227/neu.0b013e3182061ebb.

Kinoshita, Masashi, Riho Nakajima, Harumichi Shinohara, Katsuyoshi Miyashita, Shingo Tanaka, Hirokazu Okita, Mitsutoshi Nakada, and Yutaka Hayashi. 2016. “Chronic Spatial Working Memory Deficit Associated with the Superior Longitudinal Fasciculus: a Study Using Voxel-Based Lesion-Symptom Mapping and Intraoperative Direct Stimulation in Right Prefrontal Glioma Surgery”. Journal of Neurosurgery 125 (4). Journal of Neurosurgery Publishing Group (JNSPG): 1024–32. doi:10.3171/2015.10.jns1591.

Campana, Serena, Carlo Caltagirone, and Paola Marangolo. 2015. “Combining Voxel-Based Lesion-Symptom Mapping (VLSM) With A-TDCS Language Treatment: Predicting Outcome of Recovery in Nonfluent Chronic Aphasia”. Brain Stimulation 8 (4). Elsevier BV: 769–76. doi:10.1016/j.brs.2015.01.413.

Almairac, Fabien, Guillaume Herbet, Sylvie Moritz-Gasser, Nicolas Menjot de Champfleur, and Hugues Duffau. 2014. “The Left Inferior Fronto-Occipital Fasciculus Subserves Language Semantics: a Multilevel Lesion Study”. Brain Structure and Function 220 (4). Springer Nature: 1983–95. doi:10.1007/s00429-014-0773-1.

Ius, Tamara, Elsa Angelini, Michel Thiebaut de Schotten, Emmanuel Mandonnet, and Hugues Duffau. 2011. “Evidence for Potentials and Limitations of Brain Plasticity Using an Atlas of Functional Resectability of WHO Grade II Gliomas: Towards a Minimal Common Brain”. NeuroImage 56 (3). Elsevier BV: 992–1000. doi:10.1016/j.neuroimage.2011.03.022.

Brett, Matthew, Alex Leff, and John Ashburner. 2000. “Automated Nonlinear Coregistration of Damaged Brains to a Normal Template Using Cost Function Masking”. NeuroImage 11 (5). Elsevier BV: S566. doi:10.1016/s1053-8119(00)91497-6.

Ashburner, John, and Karl J. Friston. 2005. “Unified Segmentation”. NeuroImage 26 (3). Elsevier BV: 839–51. doi:10.1016/j.neuroimage.2005.02.018.

Andersen, Sarah M., Steven Z. Rapcsak, and Pélagie M. Beeson. 2010. “Cost Function Masking during Normalization of Brains with Focal Lesions: Still a Necessity?”. NeuroImage 53 (1). Elsevier BV: 78–84. doi:10.1016/j.neuroimage.2010.06.003.

Fiez, Julie A., Hanna Damasio, and Thomas J. Grabowski. 2000. “Lesion Segmentation and Manual Warping to a Reference Brain: Intra- and Interobserver Reliability”. Human Brain Mapping 9 (4). Wiley-Blackwell: 192–211. doi:10.1002/(sici)1097-0193(200004)9:4<192::aid-hbm2>3.0.co;2-y.

Werner, Rene, Matthias Wilmsy, Bastian Cheng, and Nils D. Forkert. 2016. “Beyond Cost Function Masking: RPCA-Based Non-Linear Registration in the Context of VLSM”. In 2016 International Workshop on Pattern Recognition in Neuroimaging (PRNI). Institute of Electrical and Electronics Engineers (IEEE). doi:10.1109/prni.2016.7552344.

Gordillo, Nelly, Eduard Montseny, and Pilar Sobrevilla. 2013. “State of the Art Survey on MRI Brain Tumor Segmentation”. Magnetic Resonance Imaging 31 (8). Elsevier BV: 1426–38. doi:10.1016/j.mri.2013.05.002.

Menze, Bjoern H., Andras Jakab, Stefan Bauer, Jayashree Kalpathy-Cramer, Keyvan Farahani, Justin Kirby, Yuliya Burren, et al. 2015. “The Multimodal Brain Tumor Image Segmentation Benchmark (BRATS)”. IEEE Transactions on Medical Imaging 34 (10). Institute of Electrical and Electronics Engineers (IEEE): 1993–2024. doi:10.1109/tmi.2014.2377694.

Sanjuán, Ana, Cathy J. Price, Laura Mancini, Goulven Josse, Alice Grogan, Adam K. Yamamoto, Sharon Geva, Alex P. Leff, Tarek A. Yousry, and Mohamed L. Seghier. 2013. “Automated Identification of Brain Tumors from Single MR Images Based on Segmentation with Refined Patient-Specific Priors”. Frontiers in Neuroscience 7. Frontiers Media SA. doi:10.3389/fnins.2013.00241.

Meyer, Sarah, Simon S. Kessner, Bastian Cheng, Marlene Bönstrup, Robert Schulz, Friedhelm C. Hummel, Nele De Bruyn, et al. 2016. “Voxel-Based Lesion-Symptom Mapping of Stroke Lesions Underlying Somatosensory Deficits”. NeuroImage: Clinical 10. Elsevier BV: 257–66. doi:10.1016/j.nicl.2015.12.005.

Brennan, Nicole M. Petrovich, and Andrei I. Holodny. 2016. “Use of Advanced Neuroimaging (FMRI DTI/Tractography) in the Treatment of Malignant Gliomas”. In Malignant Brain Tumors, 3–13. Springer Nature. doi:10.1007/978-3-319-49864-5_1.

Field, Aaron S., Andrew L. Alexander, Yu-Chien Wu, Khader M. Hasan, Brian Witwer, and Behnam Badie. 2004. “Diffusion Tensor Eigenvector Directional Color Imaging Patterns in the Evaluation of Cerebral White Matter Tracts Altered by Tumor”. Journal of Magnetic Resonance Imaging 20 (4). Wiley-Blackwell: 555–62. doi:10.1002/jmri.20169.

Nimsky, Christopher, Oliver Ganslandt, Peter Hastreiter, Ruopeng Wang, Thomas Benner, A Gregory Sorensen, and Rudolf Fahlbusch. 2005. “Preoperative and Intraoperative Diffusion Tensor Imaging-Based Fiber Tracking in Glioma Surgery”. Neurosurgery 56 (1). Oxford University Press (OUP): 130–38. doi:10.1227/01.neu.0000144842.18771.30.

Zhang, H, P Yushkevich, D Alexander, and J Gee. 2006. “Deformable Registration of Diffusion Tensor MR Images with Explicit Orientation Optimization”. Medical Image Analysis 10 (5). Elsevier BV: 764–85. doi:10.1016/j.media.2006.06.004.

Flavio Dell’Acqua, Marco Catani, Luis Lacerda, and Andrew Simmons. 2014. “Anisotropic Power Maps: A Diffusion Contrast to Reveal Low Anisotropy Tissues from HARDI Data.”. In Proc. Intl. Soc. Mag. Reson. Med.. Session 57. International Society for Magnetic Resonance in Medicine. http://www.ismrm.org/14/program_files/Session57.htm.

Smith, S, and T Nichols. 2009. “Threshold-Free Cluster Enhancement: Addressing Problems of Smoothing Threshold Dependence and Localisation in Cluster Inference”. NeuroImage 44 (1). Elsevier BV: 83–98. doi:10.1016/j.neuroimage.2008.03.061.

Woolrich, Mark W., Saad Jbabdi, Brian Patenaude, Michael Chappell, Salima Makni, Timothy Behrens, Christian Beckmann, Mark Jenkinson, and Stephen M. Smith. 2009. “Bayesian Analysis of Neuroimaging Data in FSL”. NeuroImage 45 (1). Elsevier BV: S173–S186. doi:10.1016/j.neuroimage.2008.10.055.

Smith, Stephen M., Mark Jenkinson, Mark W. Woolrich, Christian F. Beckmann, Timothy E.J. Behrens, Heidi Johansen-Berg, Peter R. Bannister, et al. 2004. “Advances in Functional and Structural MR Image Analysis and Implementation as FSL”. NeuroImage 23 (January). Elsevier BV: S208–S219. doi:10.1016/j.neuroimage.2004.07.051.

Jenkinson, Mark, Christian F. Beckmann, Timothy E.J. Behrens, Mark W. Woolrich, and Stephen M. Smith. 2012. “FSL”. NeuroImage 62 (2). Elsevier BV: 782–90. doi:10.1016/j.neuroimage.2011.09.015.

Berman, Jeffrey I., SungWon Chung, Pratik Mukherjee, Christopher P. Hess, Eric T. Han, and Roland G. Henry. 2008. “Probabilistic Streamline q-Ball Tractography Using the Residual Bootstrap”. NeuroImage 39 (1). Elsevier BV: 215–22. doi:10.1016/j.neuroimage.2007.08.021.

Duffau, Hugues. 2013. “Interactions Between Diffuse Low-Grade Glioma (DLGG) and Brain Plasticity”. In Diffuse Low-Grade Gliomas in Adults, 337–56. Springer Nature. doi:10.1007/978-1-4471-2213-5_22.

Jenkinson, Mark, Christian F Beckmann, Timothy EJ Behrens, Mark W Woolrich, and Stephen M Smith. 2012. “Fsl”. Neuroimage 62 (2). Elsevier: 782–90.

Leemans, Alexander, and Derek K Jones. 2009. “The B-Matrix Must Be Rotated When Correcting for Subject Motion in DTI Data”. Magnetic Resonance in Medicine 61 (6). Wiley Online Library: 1336–49.

Garyfallidis, Eleftherios, Matthew Brett, Bagrat Amirbekian, Ariel Rokem, Stefan van der Walt, Maxime Descoteaux, and Ian Nimmo-Smith and. 2014. “Dipy a Library for the Analysis of Diffusion MRI Data”. Frontiers in Neuroinformatics 8 (February). Frontiers Media SA. doi:10.3389/fninf.2014.00008.

Flavio Dell’Acqua, Marco Catani, Luis Lacerda, and Andrew Simmons. 2014. “Anisotropic Power Maps: A Diffusion Contrast to Reveal Low Anisotropy Tissues from HARDI Data”. In Joint Annual Meeting ISMRM-ESMRMB. International Society for Magnetic Resonance in Medicine.

Descoteaux, Maxime, Elaine Angelino, Shaun Fitzgibbons, and Rachid Deriche. 2007. “Regularized Fast, and Robust Analytical Q-Ball Imaging”. Magnetic Resonance in Medicine 58 (3). Wiley-Blackwell: 497–510. doi:10.1002/mrm.21277.

Smith, Stephen M. 2002. “Fast Robust Automated Brain Extraction”. Human Brain Mapping 17 (3). Wiley-Blackwell: 143–55. doi:10.1002/hbm.10062.

Keshavan, Anisha, Esha Datta, Ian McDonough, Christopher R. Madan, Kesshi Jordan, and Roland G. Henry. 2017. “Mindcontrol: A Web Application for Brain Segmentation Quality Control”. NeuroImage, March. Elsevier BV. doi:10.1016/j.neuroimage.2017.03.055.

Tristán-Vega, Antonio, Carl-Fredrik Westin, and Santiago Aja-Fernández. 2009. “Estimation of Fiber Orientation Probability Density Functions in High Angular Resolution Diffusion Imaging”. NeuroImage 47 (2). Elsevier BV: 638–50. doi:10.1016/j.neuroimage.2009.04.049.

Tristán-Vega, Antonio, and Santiago Aja-Fernández. 2010. “DWI Filtering Using Joint Information for DTI and HARDI”. Medical Image Analysis 14 (2). Elsevier BV: 205–18. doi:10.1016/j.media.2009.11.001.

Tristán-Vega, Antonio, and Carl-Fredrik Westin. 2011. “Probabilistic ODF Estimation from Reduced HARDI Data with Sparse Regularization”. In Lecture Notes in Computer Science, 182–90. Springer Nature. doi:10.1007/978-3-642-23629-7_23.

Gorgolewski, Krzysztof, Christopher D. Burns, Cindee Madison, Dav Clark, Yaroslav O. Halchenko, Michael L. Waskom, and Satrajit S. Ghosh. 2011. “Nipype: A Flexible Lightweight and Extensible Neuroimaging Data Processing Framework in Python”. Frontiers in Neuroinformatics 5. Frontiers Media SA. doi:10.3389/fninf.2011.00013.

Catani, Marco, Derek K. Jones, and Dominic H. ffytche. 2004. “Perisylvian Language Networks of the Human Brain”. Annals of Neurology 57 (1). Wiley-Blackwell: 8–16. doi:10.1002/ana.20319.

Caverzasi, Eduardo, Nico Papinutto, Bagrat Amirbekian, Mitchel S. Berger, and Roland G. Henry. 2014. “Q-Ball of Inferior Fronto-Occipital Fasciculus and Beyond”. Edited by Gaolang Gong. PLoS ONE 9 (6). Public Library of Science (PLoS): e100274. doi:10.1371/journal.pone.0100274.

Makris, N., M. G. Preti, T. Asami, P. Pelavin, B. Campbell, G. M. Papadimitriou, J. Kaiser, et al. 2012. “Human Middle Longitudinal Fascicle: Variations in Patterns of Anatomical Connections”. Brain Structure and Function 218 (4). Springer Nature: 951–68. doi:10.1007/s00429-012-0441-2.

Hofer. 2010. “Reconstruction and Dissection of the Entire Human Visual Pathway Using Diffusion Tensor MRI”. Frontiers in Neuroanatomy. Frontiers Media SA. doi:10.3389/fnana.2010.00015.

Feigl, Guenther C., Wolfgang Hiergeist, Claudia Fellner, Karl-Michael M. Schebesch, Christian Doenitz, Thomas Finkenzeller, Alexander Brawanski, and Juergen Schlaier. 2014. “Magnetic Resonance Imaging Diffusion Tensor Tractography: Evaluation of Anatomic Accuracy of Different Fiber Tracking Software Packages”. World Neurosurgery 81 (1). Elsevier BV: 144–50. doi:10.1016/j.wneu.2013.01.004.

Bernal, B., and A. Ardila. 2009. “The Role of the Arcuate Fasciculus in Conduction Aphasia”. Brain 132 (9). Oxford University Press (OUP): 2309–16. doi:10.1093/brain/awp206.

Fernández-Miranda, Juan C., Yibao Wang, Sudhir Pathak, Lucia Stefaneau, Timothy Verstynen, and Fang-Cheng Yeh. 2014. “Asymmetry Connectivity, and Segmentation of the Arcuate Fascicle in the Human Brain”. Brain Structure and Function 220 (3). Springer Nature: 1665–80. doi:10.1007/s00429-014-0751-7.

Heide, R. J. Von Der, L. M. Skipper, E. Klobusicky, and I. R. Olson. 2013. “Dissecting the Uncinate Fasciculus: Disorders Controversies and a Hypothesis”. Brain 136 (6). Oxford University Press (OUP): 1692–1707. doi:10.1093/brain/awt094.

Herbet, Guillaume, Maxime Maheu, Emanuele Costi, Gilles Lafargue, and Hugues Duffau. 2016. “Mapping Neuroplastic Potential in Brain-Damaged Patients”. Brain 139 (3). Oxford University Press (OUP): 829–44. doi:10.1093/brain/awv394.

Jezzard, Peter, and Robert S. Balaban. 1995. “Correction for Geometric Distortion in Echo Planar Images from B0 Field Variations”. Magnetic Resonance in Medicine 34 (1). Wiley-Blackwell: 65–73. doi:10.1002/mrm.1910340111.

Andersson, Jesper L.R., Stefan Skare, and John Ashburner. 2003. “How to Correct Susceptibility Distortions in Spin-Echo Echo-Planar Images: Application to Diffusion Tensor Imaging”. NeuroImage 20 (2). Elsevier BV: 870–88. doi:10.1016/s1053-8119(03)00336-7.

Pestilli, Franco, Jason D Yeatman, Ariel Rokem, Kendrick N Kay, and Brian A Wandell. 2014. “Evaluation and Statistical Inference for Human Connectomes”. Nature Methods 11 (10). Springer Nature: 1058–63. doi:10.1038/nmeth.3098.

Dixit, Prateek, and G. R. Liu. 2016. “A Review on Recent Development of Finite Element Models for Head Injury Simulations”. Archives of Computational Methods in Engineering, October. Springer Nature. doi:10.1007/s11831-016-9196-x.

Tse, Kwong Ming, Long Bin Tan, Shu Jin Lee, Siak Piang Lim, and Heow Pueh Lee. 2013. “Development and Validation of Two Subject-Specific Finite Element Models of Human Head against Three Cadaveric Experiments”. International Journal for Numerical Methods in Biomedical Engineering 30 (3). Wiley-Blackwell: 397–415. doi:10.1002/cnm.2609.

Garlapati, Revanth Reddy, Aditi Roy, Grand Roman Joldes, Adam Wittek, Ahmed Mostayed, Barry Doyle, Simon Keith Warfield, et al. 2014. “More Accurate Neuronavigation Data Provided by Biomechanical Modeling Instead of Rigid Registration”. Journal of Neurosurgery 120 (6). Journal of Neurosurgery Publishing Group (JNSPG): 1477–83. doi:10.3171/2013.12.jns131165.

Yeatman, Jason D., Robert F. Dougherty, Nathaniel J. Myall, Brian A. Wandell, and Heidi M. Feldman. 2012. “Tract Profiles of White Matter Properties: Automating Fiber-Tract Quantification”. Edited by Christian Beaulieu. PLoS ONE 7 (11). Public Library of Science (PLoS): e49790. doi:10.1371/journal.pone.0049790.

O’Donnell, Lauren J., Yannick Suter, Laura Rigolo, Pegah Kahali, Fan Zhang, Isaiah Norton, Angela Albi, et al. 2017. “Automated White Matter Fiber Tract Identification in Patients with Brain Tumors”. NeuroImage: Clinical 13. Elsevier BV: 138–53. doi:10.1016/j.nicl.2016.11.023.

Catani, M. 2005. “The Rises and Falls of Disconnection Syndromes”. Brain 128 (10). Oxford University Press (OUP): 2224–39. doi:10.1093/brain/awh622.

Duffau, Hugues. 2015. “Stimulation Mapping of White Matter Tracts to Study Brain Functional Connectivity”. Nature Reviews Neurology 11 (5). Springer Nature: 255–65. doi:10.1038/nrneurol.2015.51.

He, Biyu J., Abraham Z. Snyder, Justin L. Vincent, Adrian Epstein, Gordon L. Shulman, and Maurizio Corbetta. 2007. “Breakdown of Functional Connectivity in Frontoparietal Networks Underlies Behavioral Deficits in Spatial Neglect”. Neuron 53 (6). Elsevier BV: 905–18. doi:10.1016/j.neuron.2007.02.013.

Genova, Helen M., Venkateswaran Rajagopalan, Nancy Chiaravalloti, Allison Binder, John Deluca, and Jeannie Lengenfelder. 2014. “Facial Affect Recognition Linked to Damage in Specific White Matter Tracts in Traumatic Brain Injury”. Social Neuroscience 10 (1). Informa UK Limited: 27–34. doi:10.1080/17470919.2014.959618.

Cristofori, I., W. Zhong, A. Chau, J. Solomon, F. Krueger, and J. Grafman. 2015. “White and Gray Matter Contributions to Executive Function Recovery after Traumatic Brain Injury”. Neurology 84 (14). Ovid Technologies (Wolters Kluwer Health): 1394–1401. doi:10.1212/wnl.0000000000001446.

Duffau, Hugues. 2014. “The Huge Plastic Potential of Adult Brain and the Role of Connectomics: New Insights Provided by Serial Mappings in Glioma Surgery”. Cortex 58 (September). Elsevier BV: 325–37. doi:10.1016/j.cortex.2013.08.005.

Benzagmout, Mohammed, Peggy Gatignol, and Hugues Duffau. 2007. “Resection of World Health Organization Grade II Gliomas Involving Broca’s Area”. Neurosurgery 61 (4). Oxford University Press (OUP): 741–53. doi:10.1227/01.neu.0000298902.69473.77.

Papagno, Costanza, Marcello Gallucci, Alessandra Casarotti, Antonella Castellano, Andrea Falini, Enrica Fava, Carlo Giussani, Giorgio Carrabba, Lorenzo Bello, and Alfonso Caramazza. 2011. “Connectivity Constraints on Cortical Reorganization of Neural Circuits Involved in Object Naming”. NeuroImage 55 (3). Elsevier BV: 1306–13. doi:10.1016/j.neuroimage.2011.01.005.

